# 3D confinement physically regulates cell cycle progression in budding yeast

**DOI:** 10.1101/2025.07.16.665081

**Authors:** M Sreepadmanabh, Mridul Gautam, Nikita Bagade, Sunil Laxman, Tapomoy Bhattacharjee

## Abstract

Our understanding of cell division control traditionally comes via using liquid broths or 2D flat-plate cultures — which cannot recapitulate the complex visco-elasto-plastic properties of natural habitats such as tissues, mucus, and soil. Consequently, how such regimes of physical confinement influence proliferative growth remains unknown. Here, using engineered, mechanically tunable and transparent growth matrices, we directly visualize yeast budding and division across 3D viscoelastic regimes. We discover that elevated physical confinement drastically prolongs budding intervals without any physiological defects or activating global regulatory programs. Remarkably, extended cell division times are not associated with transcriptional or proteomic signatures of mechanosensation or cell cycle regulation. Rather, 3D confinement physically constrains the volumetric growth of incipient buds — manifesting as delayed cell cycle progression. Hence, our findings establish that physical constrainment regulates eukaryotic cell division.

## Introduction

Cell cycle progression involves a series of transitions across functionally-distinct phases, controlled by internal surveillance mechanisms ^1–4^. Molecular control over these processes is mediated by diverse regulatory elements including oscillatory cyclin-CDK expression patterns, genomic features, mitogenic factors, and metabolic modes ^5–12^. Together, this ensemble framework ensures that cellular division remains robust against systemic noise, yet sensitive to proliferation-associated mechano-chemical cues ^13–16^. However, state-of-the-art understanding of cell cycle regulation primarily derives from experiments performed using either liquid broths or 2D flat plates. This is in contrast to the natural habitats of many unicellular organisms, which are complex, disordered, and spatiotemporally heterogeneous 3D visco-elasto-plastic milieus such as tissues, mucus, and soil. Consequently, it is unclear how 3D physical confinement imposed by such environmental regimes might influence cellular division. This gap in knowledge persists because experimentally replicating the pore-scale architecture and complex shear moduli of such mechanical regimes is technically formidable ^17–21^. While past efforts have employed viscous solutions, polymeric hydrogels, and microfluidic compartments, none of these platforms capture the microporous and viscoelastic nature of granular 3D niches ^22–26^. Importantly, mechanical forces experienced by model systems such as bacterial and yeast cells in their natural habitats manifest much below experimentally-explored regimes that trigger mechanosensory pathways. Thus, it is conceivable that biophysical constraints independent of mechanotransduction may influence proliferation. We capture the essence of these ideas by asking — can purely physical constraints control a molecularly orchestrated process such as cell cycle progression?

Here, we employ the budding yeast, *S. cerevisiae* — a classical model for eukaryotic cell division studies ^27–34^ — to investigate how 3D physical confinement influences proliferative growth. We engineer optically transparent, mechanically tunable growth media with shear moduli orders of magnitude below the cellular turgor pressure, which enable single cell-level visualization of yeast budding within a viscoelastic 3D matrix. Elevating the degree of physical confinement significantly prolongs the rate of budding without causing any discernible physiological defects. Furthermore, such confinement-driven budding delays represent a transient and reversible phenomenon — manifesting solely while cells are physically embedded within the 3D growth matrix. Remarkably, cells under elevated 3D confinement do not exhibit any signatures of mechanosensory responses or differentially expressing cell cycle regulators. By systematically dissecting individual phases of the cell cycle, we find that confinement within a viscoelastic 3D matrix delays budding by physically constraining the volumetric growth of buds during the G2 phase — without affecting other phases. We argue that 3D confinement alters cell division without invoking canonical mechanosensory, metabolic, mutational, or stress-associated signaling. Rather, the physical constraints imposed by a cell’s surroundings directly restrict proliferative growth. These findings advance a model for physical confinement-driven eukaryotic cell cycle regulation, advancing our understanding of how complex mechanical microenvironments influence living matter.

## 2. Results

### 2.1. Confinement-dependent yeast growth within a 3D viscoelastic matrix

Budding yeast are widely utilized for industrial applications such as baking and brewing to ferment substrates like dough and organic matter ^35,36^. Although practical experimentations have extensively optimized for efficient yeast activity, the inability to directly visualize and model cellular processes such as growth within soft, 3D granular viscoelastic materials limit our understanding of the underlying biological phenomena. These impose conditions of 3D physical confinement on resident yeast cells, which cannot be captured using experimental platforms such as liquid broths and flat agar pads. To overcome these limitations and investigate whether the biophysical constraints imposed by 3D physical confinement influence yeast growth, we engineered soft solid-like 3D matrices formed by jammed packings of highly swollen micron-sized hydrogel granules prepared in the YPD (yeast peptone dextrose) nutrient broth (**Fig. 1A and 1B**). Each such microparticle consists of randomly cross-linked polymer chains ^37^. The swollen nature of these hydrogel granules enables jammed packings even at a low solid fraction, as well as renders the ensemble matrix optically transparent ^38^. Internally, these matrices comprise both heterogeneously-distributed micron-sized inter-particle pore spaces (θ), as well as nanometer-scale mesh sizes within each granule, which together facilitate unhindered and uniform diffusion of nutrients and small molecules throughout the bulk 3D matrix ^39^. Further characterization of the material using oscillatory shear rheology reveals a time-independent, viscoelastic, soft solid-like behavior — crucial for replicating the structural properties of environments such as dough and organic matter (**Fig. 1C**). Importantly, altering the solid fraction of the hydrogel granules allows precise control over its viscoelastic properties, enabling an interrogation of cellular growth across distinctly different mechanical regimes (**Fig. 1C, 1D, SI Fig. 1A and 1B**).

**Figure 1.**
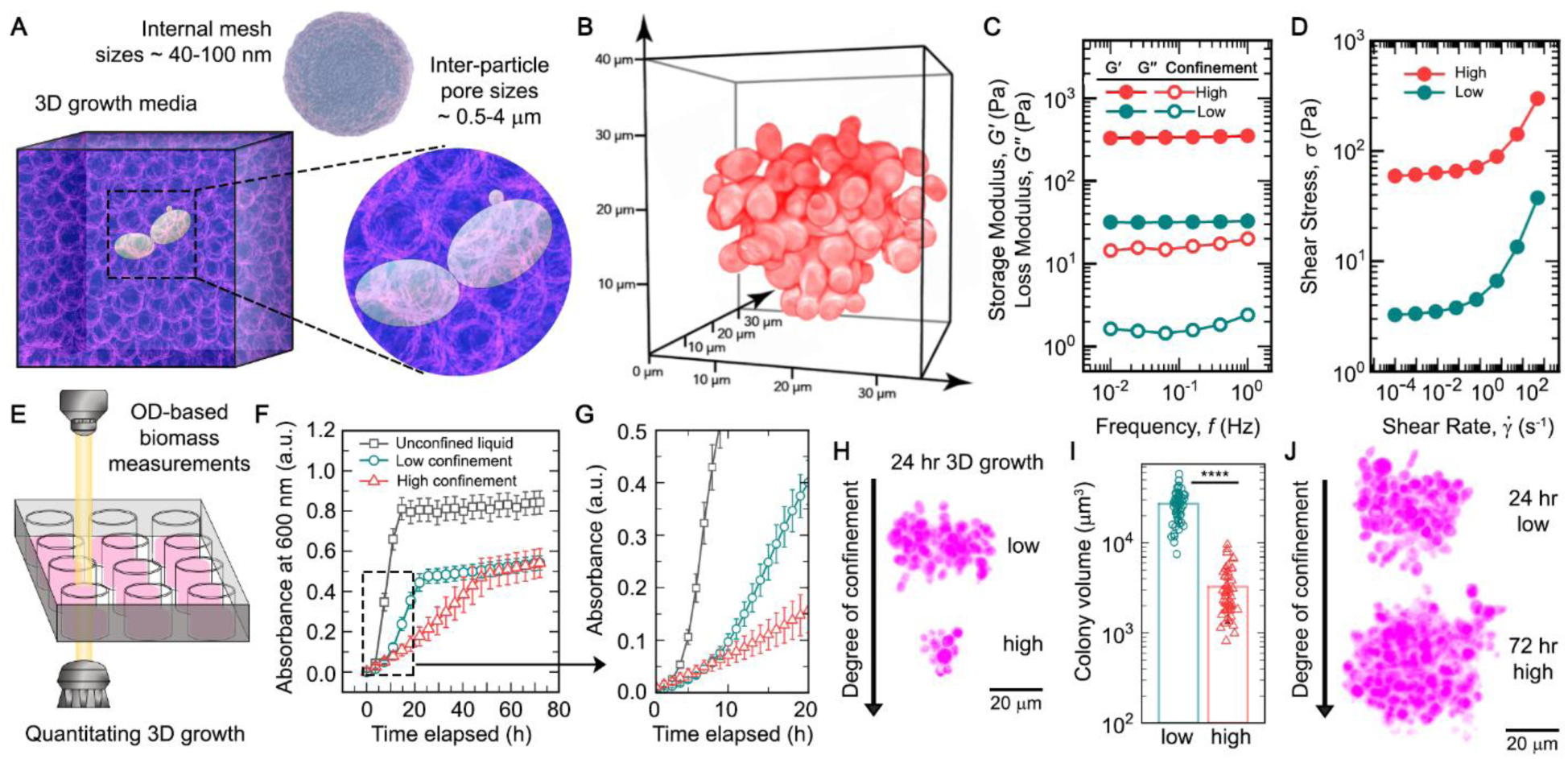
Budding yeast growth within 3D viscoelastic matrices shows a confinement-dependent decrease in proliferation. (**A**) An optically transparent, mechanically tunable matrix for visualising yeast growth within a granular viscoelastic 3D milieu. (**B**) Volumetric view of a 3D yeast colony expressing cytoplasmic mCherry (red fluorescent protein), cultured under physical confinement. (**C**) Oscillatory shear rheology measurements of the material shear moduli reveal the elastic storage modulus as being significantly greater than the viscous loss modulus, showing that the 3D growth media exhibits soft solid-like properties. (**D**) Unidirectional shear at different rates reveal a shear-dependent rheological behavior for the 3D growth media, which transitions from a fluid-like behavior under high shear regimes to a solid-like behaviour under low shear regimes. The solid-to-liquid transition occurs at the yield stress point, where both the elastic and viscous components of stress are comparable. (**E**) The transparent 3D matrix enables optical density-based absorbance measurements of the biomass production over time. (**F** and **G**) Increasing degrees of physical confinement dramatically affects the growth patterns of budding yeast. While elevated confinement decreases growth rates, the overall biomass production is comparable across both low and high confinement matrices at sufficiently long time scales. Representative data from a single biological replicate with mean +/- s.d., n = 3 technical replicates for each condition. (**H** and **I**) Colony morphometrics reveal significantly smaller colony sizes at the 24-hour timepoint in the high confinement matrices as opposed to the low confinement system. mean +/- s.d., n >= 50 individual colonies for each condition. Statistical significance calculated using an unpaired t test (∗∗∗∗ indicates p < 0.0001). (**J**) Following 72 hours of growth, colonies in the high confinement matrices appear comparable in size to those observed in the low confinement matrices at the 24-hour mark, mirroring the growth patterns suggested by optical density-based measurements. Together, the data show that elevated 3D confinement slows down growth rates without arresting the overall growth.

A key consideration is that the biophysical constraints imposed by the 3D matrix should remain invariant over the experimental time scales, without cellular growth-induced shear exerting plastic deformations on the system. We assessed this capability by subjecting the 3D matrix to unidirectional shear at different rates while measuring the shear stress response. Here, we find that the material exhibits shear-dependent rheology, by transitioning from a viscous fluid-like behavior under high shear regimes to an elastic solid-like behavior under low shear regimes (**Fig. 1D**). This transition is marked by a plateauing stress response below a threshold shear, which corresponds to the material’s characteristic yield stress (σ_*y*_) — below which the applied shear is insufficient to fluidize the system (solid-like behavior), and above which the material undergoes yielding (fluid-like behavior) ^40^. Volumetric expansion during cell growth exerts a localized shear on the surrounding matrix, which displaces hydrogel granules in its vicinity. The σ_*y*_ represents the amount of stress cells need to generate in order to rearrange the granular viscoelastic environment during growth, and hence, directly correlates with the degree of confinement imposed on cells by a given 3D growth matrix. Such rearrangements are followed by an immediate recovery once the growth-induced shear ceases, without any loss in the material properties ^41^.

By leveraging the transparency of the 3D growth matrix, we interrogated how varying degrees of physical confinement influence the growth of budding yeast. For this, we homogeneously disperse yeast cells as an initial inoculum in unconfined liquid medium, as well as both low confinement (σ_*y*_ ∼ 3 Pa and G’ ∼ 30 Pa) and high confinement (σ_*y*_ ∼ 60 Pa and G’ ∼ 330 Pa) 3D matrices. By measuring the temporal change in optical density-based absorbance values (which is proportional to the biomass increase), across all three conditions, we observe a significant decrease in growth with increasing confinement (**Fig. 1E and 1F**). Interestingly, given sufficient time to reach a steady state, the total biomass achieved remains similar across all degrees of confinement — implying that overall population-level carrying capacity is not altered (**Fig. 1F, SI Fig. 1C and 1D**). Increasing degrees of confinement slowed down the initial growth rates, indicating that elevated physical constraints within a 3D matrix merely delays proliferation (**Fig. 1G**). Notably, even an ultra-low degree of 3D confinement (σ_*y*_ ∼ 0.24 Pa and G’ ∼ 3 Pa) slows down the initial growth rate, further suggesting that the extent of decrease in growth rates is proportional to the increase in degree of confinement (**SI Fig. 1E and 1F**). This idea is further supported by quantitative measurements of volumetric colony sizes following 24 hours of growth under confinement, which show distinctly smaller colonies in the high confinement matrix (**Fig. 1H, 1I, and SI Fig. 2**). However, by the 72-hour mark, colonies formed within high confinement matrices grow sufficiently to attain sizes which are now comparable to colonies formed in the low confinement matrices at the 24-hour mark (**Fig. 1J**). Further, these characterizations also show that the observed growth rate variation is possibly due to cell-autonomous perturbations, rather than a nutrient-limitation effect due to differences in colony architecture. Indeed, the differences in growth rates manifest right from the earliest timepoints, at which stage it is unlikely that cells experience consumption-diffusion-limited nutrient deprivation. Together, these observations suggest that rather than collective effects at the population/colony level, confinement-dependent slowing of growth rates likely occurs by directly altering the budding process at a single cell level.

### 2.2. Higher confinement causes budding time delay without perturbing the cellular morphology and viability

To further dissect the confinement-dependent alteration of growth rates, we next performed time lapse imaging on single yeast cells across unconfined (liquid media), low, and high degrees of confinement (**Fig. 2A**). These results unambiguously find that an elevated degree of confinement indeed appears to slow down the budding process. To quantify this, we measured the time difference between two consecutive budding events from the same mother cell — termed as the inter-budding time (λ). As expected, we find a clear increase in the inter-budding times with increase in the degree of confinement — reiterating that confinement-induced slowing down of growth acts in a cell autonomous level by delaying the budding process (**Fig. 2B**). However, it remains unclear whether this delay is a transient response by the mother cell alone, and can be overcome in subsequent progeny cells using adaptative mechanisms. To test this, we measure the time taken by newly formed buds to fully grow and produce their own progeny cells — termed the maturation time (φ). We find that under higher confinement, maturation times of these progeny cells are also similarly prolonged, indicating that the budding time delays are not a transient response, but likely are a function of the physical environment (**Fig. 2C**). Additionally, we ensured that our sampling process is robust enough to reliably capture the distribution spread and heterogeneity inherent to single cell-level biological phenomena described using the metrics of interbudding times and maturation times (**SI Fig. 3**).

**Figure 2.**
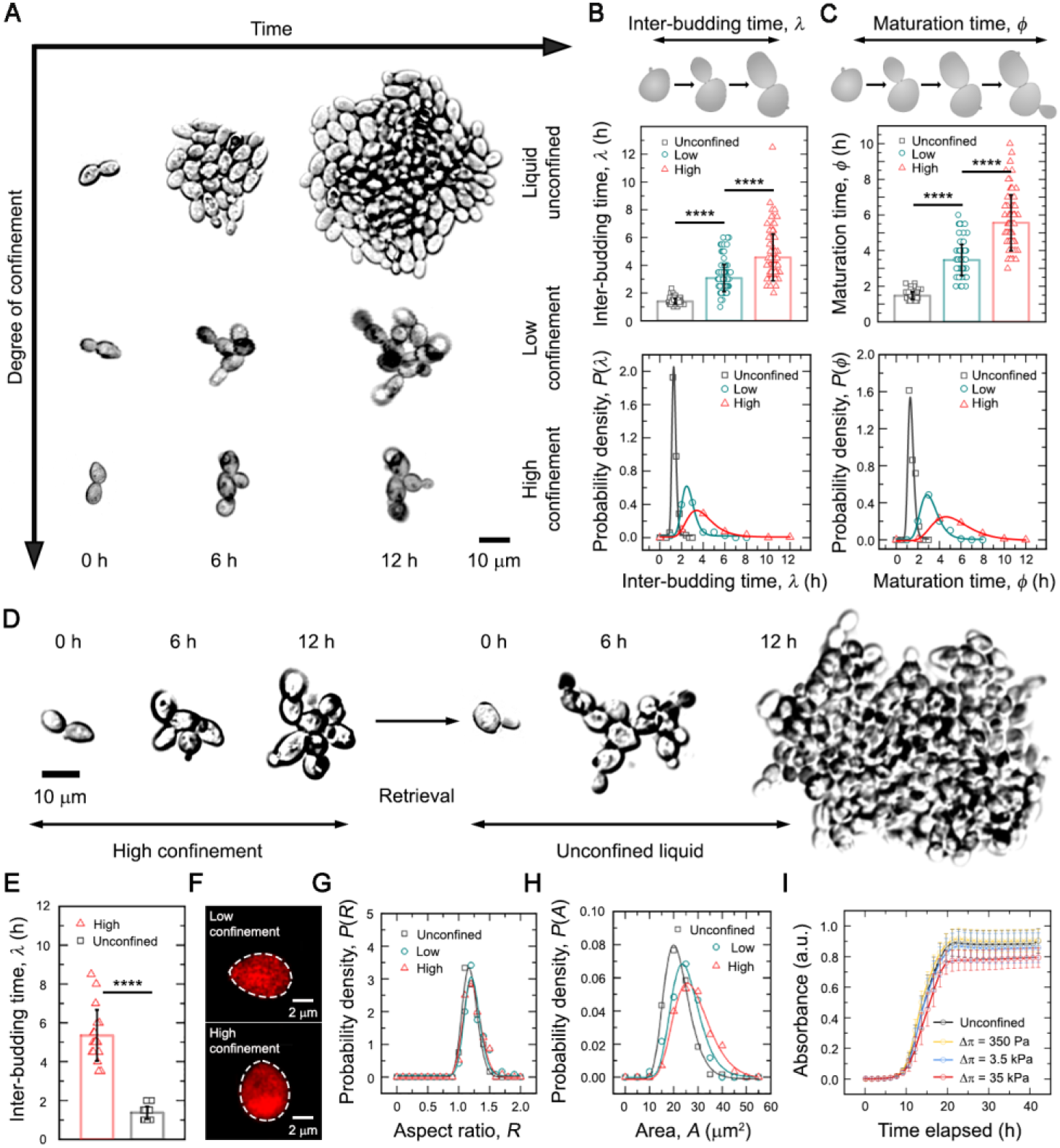
Physical confinement prolongs budding durations at the single cell level without causing physiological defects. (**A**) Growth across different mechanical regimes - unconfined liquid, low confinement 3D matrix, and high confinement 3D matrix - shows starkly contrasting effects based on the degree of confinement. Increasing confinement reduces the rate of budding-based proliferation. (**B**) Inter-budding times capture the time interval spanning one complete cell cycle, spanning two successive daughter bud emergence events from the same mother cell. Increasing degrees of confinement prolong the inter-budding times as well as lead to a more heterogeneous distribution of individual mother cell inter-budding times. mean +/- s.d., n >= 88 individual events for each condition. Statistical significance calculated using an unpaired t test (∗∗∗∗ indicates p < 0.0001). (**C**) Maturation times capture the time window spanning emergence of a daughter bud up till its own first daughter bud emergence. Similar to inter-budding times, increasing confinement severely prolongs the cell maturation times, as well as broadens the distribution of individual daughter cell maturation times. mean +/- s.d., n >= 87 individual events for each condition. Statistical significance calculated using an unpaired t test (∗∗∗∗ indicates p < 0.0001). (**D** and **E**) Confinement does not induce permanent, inheritable alterations to the cellular physiology. Cells grown in high confinement 3D media show prolonged inter-budding times, which upon retrieval and culturing in unconfined liquid media immediately revert to the wild-type phenotype, characterized by rapid budding intervals. mean +/- s.d., n = 20 individual events for each condition. Statistical significance calculated using an unpaired t test (∗∗∗∗ indicates p < 0.0001). (**F**, **G**, and **H**) Single cell morphometrics reveal no significant alterations between cells cultured in either low or high confinement matrices, suggesting that prolonged inter-budding times are not a consequence of dysmorphic morphologies. mean +/- s.d., n > 100 individual cells for each condition. (**I**) Elevation of osmotic pressure in unconfined liquid media does not significantly alter growth dynamics, even at levels 100-fold greater than the elastic modulus of the high confinement matrix. mean +/- s.d., n = 3 technical replicates for each condition.

To further ascertain whether such phenotypes emerge due to inheritable alterations in the cellular state, we retrieved cells grown in high confinement matrices and inoculated them in unconfined liquid nutrient broth. Single-cell imaging of these cross-cultured cells reveals an immediate reversion to inter-budding times consistent with those observed using cells grown entirely in unconfined liquid media (**Fig. 2D and 2E**). This data reiterates that the delay in cell division is not due to a long-term, stable adaptation process in cells, but is immediately reversible. Thus, we conclude that growth under confinement does not impose any long-term alteration in cell division times. We further assessed other attributes of cell state by visualizing the mitochondrial morphology^42,43^ and found no significant differences between cells grown in either low or high confinement matrices, suggesting that growth under confinement likely does not compromise essential metabolic functions (**SI Fig. 4**). Further, to ensure that any cell-secreted metabolic byproducts or environment remodeling factors generated during long-term growth do not modify the mechanics of the 3D growth media, or alter the degree of confinement experienced by cells over the course of these experiments, we directly measure the rheological properties of both the low and high confinement matrices (inoculated with cells) over several hours of growth at 30°C (**SI Fig. 5A**). The rheological properties remain unchanged. We also test whether cells inoculated in a medium with slightly elevated pH (∼7.4 in our 3D growth media as opposed to ∼6.5 for liquid YPD) might exhibit growth defects causative of the confinement-dependent phenotype. Towards this, we prepare liquid YPD solution with an initial pH adjusted to ∼7.4 and compare the growth dynamics of yeast cells within these against standard YPD under continuously shaken culture conditions – which again exhibit no significant differences in the growth patterns across both these regimes (**SI Fig. 5B**). We next reasoned that elevated confinement could mechanically compress the cells, thereby altering their proliferation. To check this, we perform single-cell morphology analysis of yeast grown in both low and high confinement matrices, and these analyses showed no indication of disrupted morphology (**Fig. 2F-2H, and SI Fig. 5C-5H**). Further, we also tested for potential experimental artefacts arising from the polymer formulation used to manufacture 3D growth media. We first replicated the viscoelastic properties corresponding to both low and high confinement regimes obtained using the C980 formulation with the high charge density polyelectrolyte ETD2020 ^37^ (**SI Fig. 6A and 6B**). Here, we continue to observe a confinement-dependent reduction in biomass production, which phenocopies our findings from the distinct C980-based 3D growth media (**SI Fig. 6C**).

These controls do not fully rule out osmotic stress responses under elevated confinement, which can be triggered through mechanosensory pathways and consequently contribute towards altered growth patterns ^44,45^. We therefore tested for this possibility by directly modifying the osmotic pressure exerted by unconfined liquid medium using varying concentrations of the bioinert polymer PEG-1000 to span a range extending up to 100-fold greater osmotic pressure than that exerted by our highest confinement 3D matrix (**Fig. 2I**). Remarkably, these formulations show a minimal effect on yeast growth, effectively ruling out osmotic stress responses as the underlying reason for slower growth rates. However, the 3D growth media is a granular system –– hence, a growing bud has to displace the surrounding microparticles during volumetric expansion. It is conceivable that such physical interactions could potentially trigger mechanosensory responses. In general, mechanotransduction in yeast cells is critical towards ensuring cell wall integrity and adaptability against environmental stresses. Mechanosensory responses typically involve cell wall-anchored sensory proteins, the cytoskeleton, and downstream signalling proteins –– several of which exert a profound influence on growth dynamics and cell cycle regulation ^33^. Hence, to ascertain whether confinement-dependent changes in budding dynamics are driven by mechanotransduction, we evaluated the growth of knockout strains for eight distinct, well-studied mechanosensory genes –– the primary mediator of mechanical stress-induced responses HOG1 (MAPK), the transmembrane cell wall stress sensors WSC1, WSC2, and WSC3 ^46^, the contractile ring-localizing cytoskeletal motor protein MYO1^47^, the cell wall-associated stress sensor MID2^48,49^, and the actin filament stabilizing TPM1 and TPM2 ^50,51^ –– across unconfined liquid, low confinement, and high confinement matrices (**SI Fig. 7**). These mutants all have impaired mechanotransduction based signaling. Given the wide range of functionalities covered by these genes (spanning cell wall, cytoskeleton, and mechanosensors), it is expected that these loss-of-function mutants may exhibit some heterogeneities in their growth patterns – as it is likely that the actual dynamics of cell growth and division are no longer completely identical across all eight strains. Nonetheless, we find that a loss of functionality in these mechanosensory modules did not abrogate the confinement-driven prolongation of cell cycle – rather, mutant strains also phenocopy the growth dynamics (similarly delayed division) of wild-type cells (**SI Fig. 8**). Hence, despite this diversity in their functions and sub-cellular signaling associations, we can conclude that across the board, none of these tested genes are responsible for mediating the delayed cell division response under 3D confinement. Together, these data strongly negate the possibility that mechanotransduction drives physical confinement-dependent growth patterns.

Together, these results suggest that elevated physical confinement exerts a pronounced delay on proliferative yeast growth without detrimentally or permanently altering the organismal physiology. Contrary to prior reports, we do not observe aberrant morphologies or arrested growth under 3D physical confinement, which might otherwise explain these observations ^52–55^. What, then, is the mechanistic basis underlying prolonged budding processes under high confinement?

### 2.3. Confinement delays growth without altering either the transcriptomic or proteomic landscape

Our observations thus far essentially argue that altering the mechanical environment around a yeast cell –– represented by different degrees of 3D physical confinement ––affects its proliferative growth –– represented by prolonged inter-budding time intervals. Presumably, this is indicative of delayed cell cycle progression. Historically, budding yeast represent the best-studied model system for understanding cell cycle regulation ^28–30,56^. Past efforts have also extensively profiled several environmental sensing mechanisms, many amongst which directly interface with cell cycle regulation ^57–62^. In cohesion, these modules enable yeast cells to perceive a variety of extrinsic cues and consequently respond with alterations to their proliferative growth patterns. We therefore asked if similar pathways might drive the confinement-dependent growth dynamics observed using our 3D growth media.

To better understand how elevated degrees of 3D physical confinement lead to prolonged inter-budding intervals, we first carried out a global profiling of the cellular transcriptomic state using RNA-Seq. Here, we find significantly altered transcriptional profiles between the cells grown in unconfined liquid and cells grown under confinement. This result is to be expected, and is not surprising, given the stark difference in cellular microenvironments and culturing conditions between these systems (**Fig. 3A**). Indeed, when compared against the unconfined liquid cultures, both low and high confinement-grown cells show almost a completely overlapping differential expression profile (**Fig. 3B**). In unconfined liquid cultures, yeast cells typically grow in free suspension when the culture is under constant agitation. Herein, nutrient availability remains largely uniform by virtue of advective mixing –– whereas under 3D confinement growth occurs in the form of spatially static 3D colonies, wherein nutrient molecules reach cells only via diffusion. Importantly, consumption-diffusion-dependent dynamics further impose nutrient limitations on cells buried within the interiors of large 3D colonies. This is supported by the fact that a large number of metabolism-related genes appear differentially expressed in the unconfined liquid system when compared against either of the 3D matrices (**SI Fig. 9** and **SI Fig. 10**). Together, these data strongly emphasize that unconfined liquid culturing conditions represent an entirely different biological milieu as opposed to 3D physical confinement (**Fig. 3C-3E** and **SI Fig. 10**). Hence, we suggest that cells derived from systems imposing different degrees of physical confinement must be necessarily compared only against each other –– without a direct benchmarking against unconfined liquid-grown cells –– given their fundamentally unique states.

**Figure 3.**
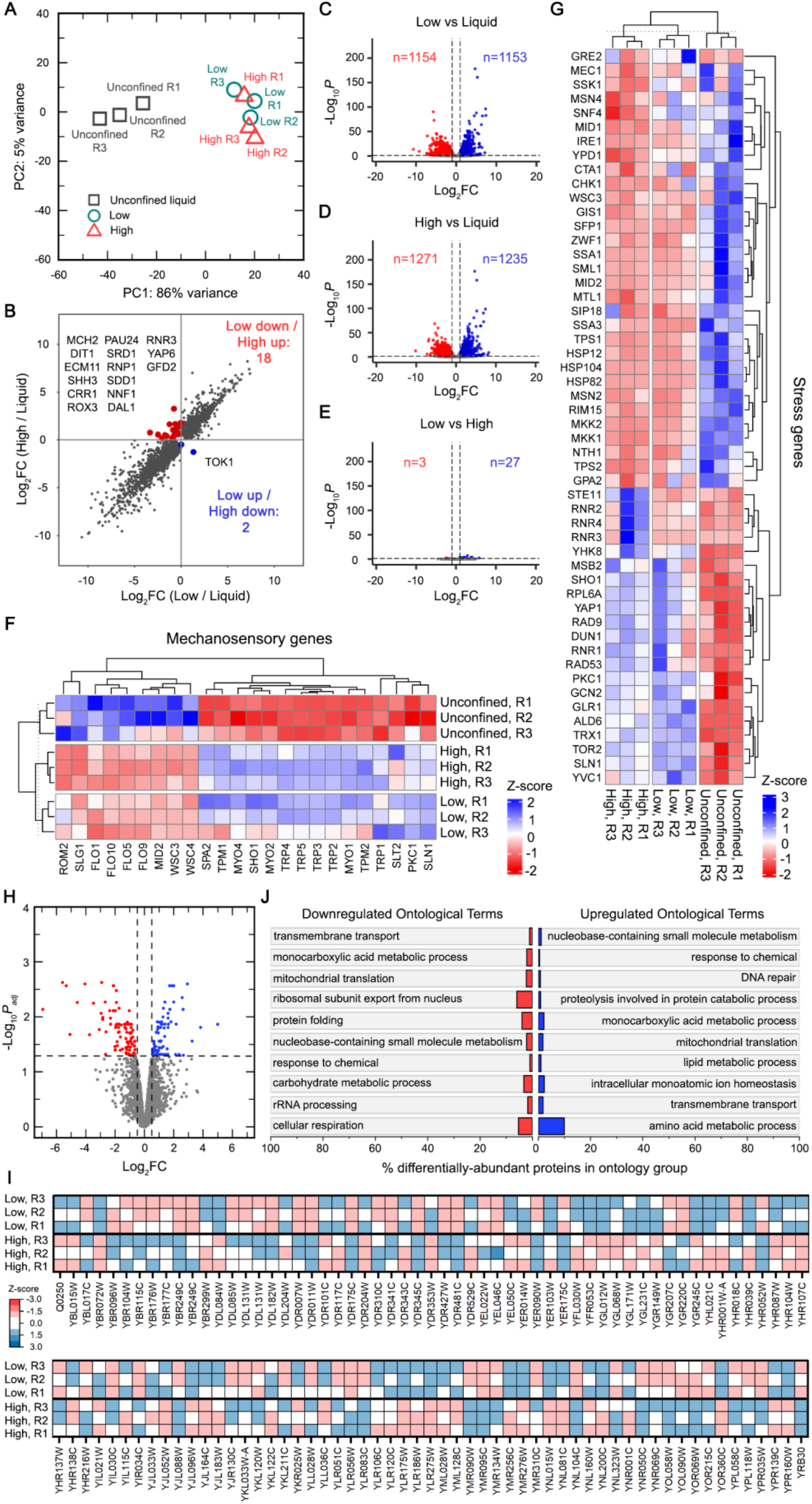
Transcriptomic and proteomic analyses of budding yeast across unconfined liquid, low confinement, and high confinement 3D growth matrices. (**A**) Principal component analyses of transcriptomic profiles for three independent biological replicates (R1-R3) across unconfined liquid, low confinement, and high confinement matrices. (**B**) Scatter plot comparing gene expression profiles between high confinement and low confinement matrices, both normalized against the unconfined liquid condition. (**C-E**) Volcano plots showing pairwise differential expression profiles across unconfined liquid, low confinement, and high confinement 3D matrices. (**F** and **G**) Heatmaps comparing the relative expression levels for genes involved in mechanosensory responses and stress responses, respectively, across all three mechanical regimes. Heatmaps show gene-wise Z-scores (row-scaled DESeq2-normalized counts) to visualize relative expression patterns across samples. Positive values (red) represent higher-than-average expression for that gene, while negative values (blue) represent lower-than-average expression. (**H**) Volcano plot showing differentially abundant proteins across cells grown under either low or high confinement, with thresholds set as adjusted p-value < 0.05 and log2FC >= |0.5|. (**I**) Abundance values rescaled as a z-score for all the annotated differentially abundant proteins obtained from quantitative proteomics analysis, compared as high confinement vs low confinement. (**J**) Ontological classifications of biological processes being either downregulated or upregulated between cells grown under high confinement vs low confinement 3D matrices, represented as a percentage of the total number of members in each such grouping.

Quite remarkably, however, we observe almost no meaningful differential expression between cells in high confinement matrices compared against the cells in low confinement matrices (**Fig. 3E**). Only a handful of statistically significantly differentially expressed genes are observed between low and high confinement systems (**Table S1**). Among these, we do not find any representation from canonical mechanosensory (**Fig. 3F**) or stress responses-associated effectors (**Fig. 3G**) –– strongly suggesting that an increase in the degree of confinement does not induce any significant transcriptional response associated with these pathways. It should be noted that canonical mechanosensory proteins such as MYO1 and TPM2, among others, only exhibit differential regulation when compared between liquid-grown and 3D confinement-grown cells. This is likely a consequence of the differences arising from growth within an unconfined liquid-like regime as opposed to a 3D mechanical regime. However, when comparing between cells grown across the low and high confinement matrices, we do not observe any such differential expression associated with mechanosensory responses – suggesting that an increase in the degree of confinement within these regimes does not trigger canonical mechanotransduction.

It is conceivable that despite similarities at the transcriptional level, biochemical signaling involved in mechanosensory or stress-associated responses could be differentially regulated at the protein level. To better explore this possibility, we performed a quantitative proteomics analysis between slow-dividing cells cultured in high confinement 3D matrices and fast-dividing cells cultured in low confinement 3D matrices following 24 hours of growth. Remarkably, these results were in excellent agreement with our transcriptomic analyses – the proteomics data demonstrated negligible differences between high confinement and low confinement-grown cells (**Fig. 3H** and **3I**). Further, we find no signatures of altered mechanosensation or stress responses based on proteomic changes. The limited number of proteins identified as differentially abundant is in stark contrast to the hours-long delay in cell cycle progression in high confinement matrices, which would appear likely to manifest a much larger footprint of altered regulatory programs (**Table S2**). In fact, there is almost no alteration of any major signaling network, implying that cell states under high confinement remain in very similar states to cell states under low confinement (**Fig. 3J**). Despite the highly limited effect of these differential protein abundances on altering biological functionality (as evidenced by the very low coverage of all top-ranked ontology groups, (**Table S3**)), it is worthwhile to discuss some broad differences. Across both downregulated and upregulated ontologies, there is a marked representation of metabolism-associated processes. This is readily explained by the fact that proteomic analyses are performed using samples harvested following 24 hours of growth under 3D confinement, at which point, the differences in sizes between larger colonies in low confinement and smaller colonies in high confinement (**Fig. 1H, 1I,** and **SI Fig. 2**) imply that different proportions of the populations in both are nutrient-limited due to their positioning within the core of the colony, and hence, these cells likely exhibit differences in the metabolic states. This is merely a consequence of the limitations imposed by the competition between consumption-diffusion dynamics, whereby, cells at the periphery of the colony enjoy sustained nutrient access, whereas, cells buried in the colony’s core are likely to be nutrient-limited. However, this does not imply that metabolism is the key driver for differences in inter-budding rates, as cells at all stages of growth - from early-stage, isolated, and well-fed single cells to late-stage colonies of cells - exhibit the same type of confinement-dependent prolongation of budding intervals. If indeed such a metabolic influence were to exist, we would have observed a progressive shift in the patterns of confinement-dependent prolongation of budding between fast-budding cells under low confinement and slow-budding cells under high confinement - however, there is no such temporal adaptive response observed in any of the experiments, either at the bulk population level (growth curves) or single cell-level (time lapse microscopy). Furthermore, prior work has extensively demonstrated the effects of nutrient starvation on cell cycle arrest / delay (**SI Fig. 14**) - and all of these studies have demonstrated that such a mode of action induces large-scale changes in gene expression levels, none of which are observed in our case. Rather, the physical confinement-dependent prolongation of budding intervals is distinguished by its vanishingly small number of differentially-expressed genes / differentially-abundant proteins, which do not strongly correlate with any major change in the cell’s biological functionality that would be likely to either disrupt cell cycle regulation, or trigger stress responses, metabolic shifts, and mechanosensation.

While the results from both transcriptomic and proteomic profiling substantiate the idea that confinement-induced budding time prolongation does not trigger a detectable level of differential gene expression or differential protein abundance between fast-budding cells under low confinement and slow-budding cells under high confinement, it is necessary to consider a couple caveats. First, the 24-hr timepoint of interrogation employed here for the omics-based analyses do not entirely rule out the possibility of early/transient signaling responses, which may arise during the initial phases of growth. However, it is also worthwhile to note that we do not observe any form of cellular adaptation to confinement over time. Across both low and high confinement systems, budding events captured at both the early stage (first few hours) and late stage (several hours post-inoculation) do not exhibit any significant deviations. Rather, both originally inoculated mother cells, and their daughter cells (born and matured hours later) continue to follow the overall trend of either fast budding under low confinement or slow budding under high confinement. We also note that in all the absorbance-based growth assays, we do not observe any aberrant shift in the slopes of the curves corresponding to either low or high confinement-grown cells – suggesting that there is no significant difference in the growth modes between short and long time scales. Hence, even if early transient responses do exist, they likely do not contribute significantly towards the confinement-dependent delay of cell cycle progression as reported herein. Second, the techniques of bulk RNA-Seq and quantitative proteomics as employed here cannot capture the single-cell / sub-population-level heterogeneity in cellular states. In fact, this constitutes a very interesting problem with regard to spatially structured environments such as the 3D growth matrices presented in this work, as the imposition of physical constraints is likely to amplify sub-population-level heterogeneities. However, a systematic investigation of such effects would require sophisticated techniques such as spatially-resolved single cell / single colony retrieval and sequencing, which, unfortunately, lie beyond the scope of the present study. However, from both a fundamental biology and bioengineering perspective, dissecting such granular differences within clonally-identical populations in complex environments represents a very exciting domain of enquiry, with broad implications for understanding the evolution of spatial structuring in natural habitats.

The key implication here is that confinement-induced budding time prolongation — in itself a drastic alteration of the intricately orchestrated cell cycle process — manifests without requiring any major changes to the transcriptional or proteomic landscape. Considering this, we now hypothesize that 3D confinement leads to delayed cell cycle progression not via mechanosensory responses or disrupted proliferative signaling, but rather, via a physical constriction of the budding process by the 3D matrix.

### 2.4. Confinement targets volumetric growth during G2 phase of the cell cycle which delays the overall yeast growth

To understand how growth under 3D confinement regulates the cell cycle, we dissect the kinetics of cell cycle progression in budding yeast (**Fig. 4A** and **SI Fig. 11**). We employed genetically modified lines expressing fluorescently tagged proteins involved in cell cycle regulation — CDC14 and CDC10 (**Fig. 4A**) — which grow similar to wild-type cells under unconfined liquid conditions (**SI Fig. 12A**). We first considered the CDC14, which is a phosphatase involved in mitosis regulation by counteracting CDK activity through dephosphorylation — thereby, acting as a prerequisite for mitotic exit and subsequent entry into G1 phase ^63–65^. The fusion reporter form of this exhibits a puncta-like signal in the mother cell, which subsequently splits into the daughter bud. This event marks the anaphase during late M phase — hence, the time interval between each successive puncta-splitting event (the inter-punctating time, ρ) approximates the overall cell cycle duration (**Fig. 4B** and **4C**). Quantitative analyses of timelapse imagings performed on cells in both low and high confinement matrices show a strong confinement-dependent effect, wherein, cells from high confinement matrices exhibit significantly prolonged inter-punctating times (**Fig. 4D**). To control for the possibility that the addition of a fluorescent tag to a key cell division protein may alter the cell division kinetics, we also measure inter-budding times for the CDC14-mNeonGreen reporter cells. Despite minor variations in the exact values of interbudding times between wild-type and reporter cells (likely an inherent limitation due to the fluorescent tag on CDC14), we confirm that the central phenotype of prolonged interbudding times under higher confinement remains well-conserved and in good agreement between both wild-type and reporter strains (**Fig. 4E**). Moreover, we also found an excellent agreement between inter-budding times and the CDC14 inter-punctating times (**4F**, and **SI Fig. 12B**). Interestingly, the duration of anaphase — indicated by the puncta splitting time interval (ψ) wherein a parent punctum splits into two low-intensity dispersed patches before forming two well-separated, condensed puncta — remains largely invariant across both degrees of confinement (**Fig. 4G** and **SI Fig. 12C**). It should be noted that these analyses do not specifically discriminate between initially-inoculated mother cells and their progeny that go on to produce daughter buds. Rather, all specimen demonstrating a complete progression of cell cycle were considered for these analyses. Together, these data strongly support the hypothesis that physical confinement leads to an overall extension of the cell cycle duration under higher confinement, as expected from the delayed budding times. However, this does not rule out the possibility that cells could be undergoing extended cycle progressions due to arrest in G0 phases.

**Figure 4.**
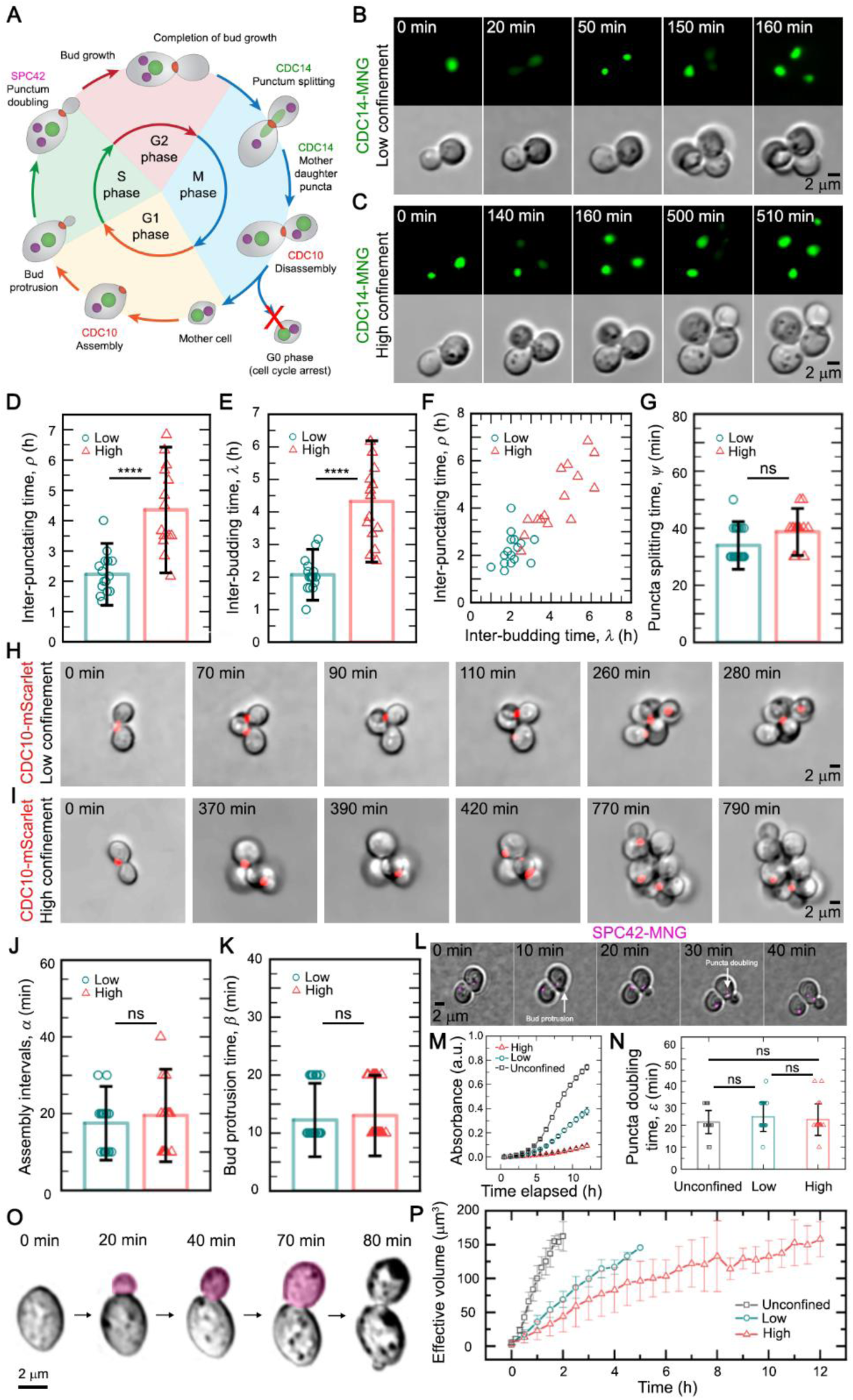
Biophysical dissection of cell cycle progression. (**A**) Schematic representation of cell cycle progression in budding yeast, highlighting the specific phase-based patterns of CDC14, CDC10, and SPC42 activity. (**B**-**G**) Tracking of CDC14-mNeonGreen (MNG) localization dynamics to quantify cell cycle progression. (**B** and **C**) Localized CDC14-mNeonGreen punctum in the mother cell split into diffused patches during the anaphase, followed by condensation into spatially separate puncta in both mother and daughter cell following completion of M phase, marking the cell cycle end. (**D**, **E,** and **F**) The inter-punctating time intervals for a given mother cell are severely elongated for cells in the high confinement matrix as opposed to cell in the low confinement matrix, which also correlates with the prolonged inter-budding times. mean +/- s.d., n >= 15 individual events for each condition. Statistical significance calculated using an unpaired t test (∗∗∗∗ indicates p < 0.0001). (**G**) However, the duration for which a CDC14-mNeonGreen puncta remains diffused during splitting in the anaphase is largely similar for low and high confinement-cultured cells, indicating that this phase of the cell cycle is not affected by elevated confinement. mean +/- s.d., n >= 23 individual events for each condition. Statistical significance calculated using an unpaired t test (ns indicates non-significant for p = 0.0021, since the absolute difference between the means of low and high confinement datasets is less than half the duration of a single imaging interval (10 minutes)). (**H-K)** Tracking of CDC10-mScarlet localization dynamics to quantify cell cycle progression. (**H** and **I**) CDC10-mScarlet localizes to the bud site prior to bud protrusion in the G1 phase and remains associated with the bud neck region until (**J**) The time interval between disassembly and assembly of the CDC10-mScarlet marks the time period between completion of the cell cycle and commitment to initiating the next cycle. This does not significantly differ between either low or high confinement-grown cells, indicating that no differential G0 arrest occurs, nor is the early G1 phase delayed under confinement. mean +/- s.d., n >= 20 individual events for each condition. Statistical significance calculated using an unpaired t test (ns indicates non-significant for p > 0.01). (**K**) The period between assembly of CDC10-mScarlet at the bud site and emergence (protrusion) of the initial bud at the start of S phase is also not significantly different between cells in low and high confinement systems. Together, these observations using CDC14-mNeonGreen and CDC10-mScarlet suggest that neither the G1 nor late M phase are significantly delayed under elevated confinement, nor do the cells get differentially arrested in a G0 phase. mean +/- s.d., n >= 40 individual events for each condition. Statistical significance calculated using an unpaired t test (ns indicates non-significant for p > 0.01). (**L**) Timelapse imaging showing the spindle pole body marker (SPC42-mNeonGreen), pseudo-colored in magenta for contrast, overlaid on the brightfield micrograph. The start of the S phase is marked by the earliest instance of bud protrusion, whereas, conclusion of the S phase is indicated by a duplication of the SPC42-MNG puncta. (**M**) Optical density-based absorbance measurements of biomass production for the SPC42-MNG strain used in these experiments, across unconfined liquid as well as both low and high confinement 3D growth media, showing that increase in confinement delays cell cycle progression. (**N**) Direct measurements of the S phase duration using single-cell microscopy, demonstrating that the S phase duration is nearly identical across all three mechanical regimes, and validating that the S phase duration remains conserved despite the overall differentially-prolonged cell cycle under confinement. mean +/- s.d., n >= 20 individual events for each condition. Statistical significance calculated using an unpaired t test (ns indicates non-significant for p > 0.01). (**O**) Schematic illustration of quantitative single-cell measurements for bud growth. (**P**) The rate of increase in bud size shows a strong dependence on the degree of confinement, mirroring the patterns observed for inter-budding times across different mechanical milieus. Cells in unconfined liquid show rapid increase in bud size, whereas cells under 3D confinement exhibit a much slower bud growth — with cells under low confinement showing faster growth than cells under high confinement. mean +/- s.d., n >= 20 individual events for each condition.

Next, we considered the CDC10, a core structural component of the septin ring, which is essential for the assembly and stability of septin at the presumptive bud site just prior to bud protrusion in G1 phase ^66–69^. Upon the completion of cytokinesis (septation), marking the end of cell cycle, CDC10 undergoes disassembly, followed by re-assembly at the next presumptive bud site (**Fig. 4H and 4I**). Notably, we observe that despite the inter-budding times being prolonged under elevated confinement (**SI Fig. 12D and 12E**), the time interval (α) between the late M phase –– marked by CDC10 disassembly –– and the early G1 phase –– marked by CDC10 assembly at the new bud site –– remains conserved across each of unconfined liquid, low confinement, and high confinement matrices (**Fig. 4J and SI Fig. 12F**). This suggests that the prolongation of cell cycle under high confinement cannot be attributed to differentially induced G0 phase arrest under confinement (**Fig. 4A**). Additionally, CDC10 assembly marks the Start checkpoint, at which stage cells commit to budding. This culminates with a bud protrusion event, marking the beginning of the S phase ^70,71^. Interestingly, the time interval between CDC10 assembly and bud protrusion (β) also remains similar across each of unconfined liquid, low confinement, and high confinement matrices (**Fig. 4K and SI Fig. 12G**). Together, these results confirm that the overall G1 phase duration remains conserved regardless of the external mechanical milieu –– indicating that cells across all three mechanical regimes proceed up till the S phase without differentially experiencing any environment-induced delays.

We next considered the possibility that the S phase could be aberrantly elongated under high confinement. However, the large difference in inter-budding times between low and high confinement would necessitate an hours-long prolongation of the S phase. Prior reports have extensively characterized stress markers corresponding to arrested replication^72–78^. Similar signatures being differentially upregulated for cells under high confinement would point towards replicative stress arising from a prolonged S phase. Both our transcriptomic and proteomic analyses discredit this possibility, as we do not observe any signatures of either DNA damage or replicative-stress response pathways being differentially upregulated due to elevated physical confinement (transcriptomics – **Fig. 3G, SI Fig. 9, SI Fig. 13,** and **Table S1;** proteomics – **Fig. 3J** and **Table S2**). Alteration of DNA replication-associated signaling is only observed when unconfined liquid-grown cells are compared against either low or high confinement 3D matrices-grown cells. However, this can be directly attributed to the differences in physical milieus, going from a liquid-like environment to 3D confinement. Such liquid-to-3D transitions also trigger significant alterations to the metabolic programs, that may in turn crosstalk with the replication-associated regulatory pathways. Interestingly, no such differences are observed when cells from low confinement matrices are compared against high confinement matrices, despite the large differences in their cell division times. These observations are supported by the ontology classifications for both RNA-Seq and quantitative proteomics data (transcriptomics –**SI Fig. 9;** proteomics – **Fig. 3J** and **Table S3**), wherein, DNA replication stress is not identified as an affected process when compared across cells grown under high vs low confinement matrices. Given the conserved nature of replication stress responses as well as the multiple signalling safeguards employed by cells to detect such aberrations, the lack of any such signatures strongly supports the idea that S phase is not the differentially elongated regime contributing towards prolonged cell cycle duration for cells in the high confinement matrices.

Additionally, using a fluorescently tagged construct for the spindle pole body marker SPC42, we directly measure the duration of the replicative S phase. The fusion reporter form of this (SPC42-mNeonGreen) exhibits a puncta-like signal in the mother cell, which subsequently multiples into two separate puncta following S phase replication. Here, we track the time period between bud protrusion (indicative of S phase commencement) and clearly discernible duplication of the SPC42-MNG puncta (indicative of S phase completion) – which gives an estimate for the duration of the S phase (**Fig. 4L**). As a control, we verify that this reporter strain also exhibits confinement-dependent reduction in growth rates, similar to the wild-type cells (**Fig. 4M**). From our single-cell microscopy-based measurements we quantitively show that the duration of the S phase remains highly conserved across both low confinement and high confinement 3D matrices, as well as the unconfined liquid cultures (**Fig. 4N**). Therefore, these analyses allow us to eliminate the S phase as being aberrantly prolonged, leaving only volumetric bud growth during the G2 phase as the likely target for confinement-driven prolongation.

Since G2 is where the budding cell body undergoes volumetric growth, we assess the change in cell size with time by measuring the growing bud’s radius right from its emergence until the successive budding event (bud protrusion) from its mother cell (**Fig. 4O**). Measurements of budding cell sizes across time reveals that while cells in both low and high confinement matrices achieve a comparable maximum size (also demonstrated in **Fig. 2F-H**, **SI Fig. 5B-5G,** and **SI Fig. 12H**), there occurs a pronounced time delay in the high confinement matrices (**Fig. 4P**). Indeed, as the degree of confinement increases, the rate of volumetric growth concomitantly decreases. These effects are likely a consequence of mechanical constraints on volumetric growth during the G2 phase by the high confinement matrix, forcing budding yeast cells to generate higher shear in order to locally rearrange the surrounding granular matrix and reach the threshold size required to initiate mitosis. It is useful to present these findings against the classical sizer-timer model^79–81^ of cell cycle regulation in budding yeast, which posits that size-based thresholds regulate passage through Start, followed by temporally-restricted subsequent phases^82^. While this model provides valuable insights on the modalities of cell cycle regulation, recent work has unearthed additional control strategies – most pertinently, that size-based regulation can be exerted even during G2 and M phases via nutrient signalling and cytoskeletal perturbations^83–86^. Similarly, our work demonstrates a mode of cell cycle regulation via physical constriction of G2 volumetric growth under 3D confinement. Such confinement-dependent effects do not negate the fact that both G1 sizer and S-G2-M timer controls critically regulate cell cycle progression. Rather, our results enhance the existing paradigm by demonstrating a physical confinement-driven model which putatively operates alongside the genetically-encoded sizer-timer regulatory framework.

## 3. Discussion

Our work establishes 3D physical confinement as a fundamentally unique regulator of eukaryotic cell division, without invoking mechanosensory, metabolic, mutational, or stress-associated signaling pathways (**SI Fig. 14**)^87–91^. Here, we accomplish microscopic visualization of yeast budding within a viscoelastic granular environment using optically transparent 3D growth media. Elevated physical confinement within these matrices severely prolongs budding — without resulting in any growth defects or long-term alterations. Quite remarkably, even an hours-long increase in budding time intervals does not manifest either transcriptomic or proteomic signatures associated with mechanosensation or stress signaling. Rather, the volumetric growth of incipient buds is physically constrained by its environment, resulting in delayed cell cycle progression. Presently, cell cycle regulation is conceptualized as a multi-scale framework encompassing myriad elements such as protein complexes (cyclins and CDKs), oscillatory gene expression programs, epigenetic landmarks, metabolic activity, chemical inputs (mitogens), mechanosensory cues (stiffness and osmotic pressure), as well as the precursor cell state ^28–30^. However, a near-universal approach is to ultimately ascribe specific subsets of molecular agents as the mechanistic effectors cognate to observed phenomena. In contrast, our findings advocate that the physical microenvironment of a cell can exert significant regulatory influence on proliferative growth without any discernible changes in terms of differential gene expression levels or differential protein abundance. Specifically, we do not detect any transcriptional/protein-level signatures suggestive of selective activity corresponding to canonically-identified mechanosensory, metabolic, mutational, or stress-associated signaling pathways – implying that the differential activation of any such pathways leading to altered downstream signaling is likely not the reason why physical confinement delays cell cycle progression. However, it is also important to acknowledge that the present study does not rule out the prospect of regulatory effects mediated by mechanisms such as altered protein localization, selective post-translational modifications, shift in phosphorylation states, changes to sub-cellular trafficking, or disrupted polarity establishment.

While in principle these constitute valid possibilities, it is beyond the scope of this first paper to test for all such scenarios, given both the technical complexity and massive number of potential candidates. However, we anticipate that the foundational results presented in this work will nucleate a broader field of enquiry. Importantly, the paradigm we establish in this work holds significant implications towards exploring how proliferative growth of eukaryotic cells inhabiting mechanically complex natural environments such as soil, mucus, and tissues are governed by the spatiotemporally dynamic material properties of their surroundings.

It is interesting to contrast the 3D physical confinement-driven delay in cell cycle progression for budding yeast against prior work imposing compressive stresses on yeast cells^92,93^ While delays/arrests in cell cycle progression have been reported, it is important to distinguish that prior work employs rigid confinement within microfluidic setups, generating compressive stresses of ∼400 kPa, as well as imposes a highly crowded condition on the cells, which may itself induce nutrient deprivation-associated stress responses. By contrast, the degrees of physical confinement explored in this study operate at least three orders of magnitude below these limits, with the highest degree of confinement being a G’ ∼ 350 Pa (0.35 kPa). Furthermore, the viscoelastic systems in this study lend themselves to elastic deformations under growth-generated pressures, enabling localized rearrangements of the microparticles to create space for newly-generated cells. Hence, while the effect of mechanical shear and compressive stresses have been traditionally explored at scales much higher than those employed in this work, we speculate that the attribution of specific mechanosensory pathways as the underlying mechanisms may in part derive from the high magnitude of stress. Under regimes much closer in stiffness to soft viscoelastic media such as mucus and tissues (∼<1 kPa), the canonically-identified mechanosensory pathways are likely to remain inactive, due to the external mechanical inputs falling far below their sensing threshold. This opens up the possibility for physical constraints-based reduction of volumetric growth rates to manifest their effects without requiring any alterations to either gene or protein expression profiles. In addition to biophysical regulation of cell cycle delay, there still remains a unique possibility that there exist an as-yet-unknown class of mechanosensory proteins which possess a tightly-controlled sensing window active only under ultra-low degrees of confinement and do not trigger major gene-expression programs. This presents a rich set of questions for future investigations aimed at decoding how cellular sensing operates at such ultra-low mechanical stress regimes.

A complementary aspect of proliferative growth — cellular aging — has been extensively studied using budding yeast as a model system. Herein, two broadly distinct paradigms of aging prevail: replicative lifespans (RLS) and chronological lifespans (CLS)^94^. RLS is defined by cell cycle-centric division processes up until senescence, whereas CLS captures the post-mitotic survival time of non-dividing cells. Present understanding identifies the G0 phase as a stage bridging proliferative (RLS) and quiescent (CLS) states; with the induction of G0 itself being primarily attributed to nutrient deprivation, oxidative and osmotic stresses, as well as acidic microenvironments^95^. Our work here offers a parallel perspective on these dynamics. By prolonging the cell cycle without causing quiescence-induced arrest, higher confinement presumably pushes cells into a hitherto uncharted regime which cannot be fully captured using the conventional definitions of RLS and CLS paradigms. It is hence interesting to ask — what do such altered proliferative timespans imply for the aging process? Do slow-cycling cells age slower due to reduced genomic divisions, or, do they compromise fitness due to increased accumulation of deleterious mutations? Our platform offers a unique opportunity to dissect these questions by offering a tunable physical control over cell cycle durations without necessitating chemical, genomic, or metabolic perturbations. Future work along these lines could potentially redefine how we understand and classify the processes of cellular aging and senescence, particularly with regard to growth within complex 3D microenvironments.

## 4. Materials and Methods

### 4.1. Preparation of 3D growth media

We prepare 3D growth media by homogeneously dispersing lyophilized granules of either Carbopol C-980 (Ashland) or Carbopol ETD2020 in liquid Yeast Peptone Dextrose (YPD) broth at concentrations between 0.4% (ultra-low confinement), 0.5% (low confinement), and 1% (high confinement for C-980) / 1.1% (high confinement for ETD2020) – all values given are in *w/v%*. Following complete dissolution, we neutralize the pH of the suspension to pH ∼ 7.4 using 10N NaOH – as the C980 and ETD2020 consists of negatively charged randomly crosslinked polymer chains. This causes the hydrogel granules to swell up and form a jammed packing, resulting in a disordered, granular, and porous 3D matrix. Due to the low solid fraction and highly swollen state of individual granules, the ensemble material is rendered optically transparent.

### 4.2. Rheological characterization

We characterize the rheological properties of these 3D packings using a shear-controlled Anton Parr (MCR302e) rheometer with a roughened cone-plate geometry. By subjecting 3D growth media samples to unidirectional shear at varying rates, we record the consequent shear stress responses as a function of the shear rates. This reveals a shear-dependent rheology, wherein the material behaves like a fluid under high shear rates while reversibly transitioning to a soft solid-like behavior under lower shear rates. The transition point between these two regimes is captured by the crossover shear rate, which represents the shear rate at which both the viscous and elastic components of the shear stress are equal. This directly corresponds to the material’s characteristic yield stress, which we employ as a measure of the degree of confinement. To further characterize the viscoelastic material properties of these 3D growth media, we apply small amplitude (1%) shear at varying frequencies and record the complex shear moduli – which is further resolved into components of elastic storage modulus (G’) – which represents the energy stored during deformation, and thereby, the elastic solid-like nature – and viscous loss modulus (G”) – which represents the energy lost as heat and frictional losses during deformation, and thereby, the viscous fluid-like nature. We observe that under low shear regimes, the jammed microgel formulations behave as a soft-solid – characterized by G’ > G” – which also remains invariant of the frequency at which the shear is applied, indicative of time-independent material properties^96^. Hence, our 3D growth media provide a stable, self-healing, soft solid-like platform for culturing yeast cells under tunable degrees of physical confinement. As a comparison, we have contrasted the material properties of the 3D growth media used in this work against prior measurements of mucosal samples (**Table S4**)^39,97–105^. For measuring the rheological properties of the 3D microgel media over several hours of growth, we perform the above steps while maintaining the sample loading stage at 30°C.

### 4.3. Quantifying porosity of the 3D growth media

To quantify the porosity of the jammed microgels, we track the thermal diffusion-guided motion of 200nm tracer fluorescent beads homogeneously dispersed throughout the 3D matrix. Timelapse imaging of single bead diffusions are acquired using point-scanning laser confocal microscope (Nikon AIR HD25). Using a custom-written MATLAB script based on the Crocker-Grier algorithm, we track individual particle trajectories with sub-pixel precision to obtain the mean-square displacements (MSD) as a function of lag times. Over short time scales, beads freely explore the inter-particle pore spaces, indicated by a linear increase in MSD with lag time. However, over longer time windows, their motion becomes constrained by the pore boundaries, indicating by plateauing MSD values. We identify such plateaus – the square root of which added to the particle diameter gives an estimate for the inter-particle pore space dimensions explored by the bead. Such measurements are performed for approximately fifty beads in both low and high confinement regimes, and we derive population-level statistics for the matrix porosity by representing pore sizes as a 1-CDF (cumulative distribution function).

### 4.4. Yeast culture in liquid broth and 3D growth matrix

In this work, we employ the prototrophic budding yeast (*S. cerevisiae*) strain CEN.PK MATa. Additionally, the following strains have been used – *S. cerevisiae* genetically modified to constitutively express either cytoplasmic red-fluorescent protein mCherry or mitochondria-localizing green-fluorescent protein mNeon-Green; genetically engineered *S. cerevisiae* carrying fluorescent fusion constructs which tag CDC14 with the green fluorescent protein mNeonGreen and CDC10 with the red fluorescent protein mScarlet; genetically engineered *S. cerevisiae* carrying a fluorescent fusion construct which tags SPC42 with the green fluorescent protein mNeonGreen; and eight different strains of *S. cerevisiae* which have a genomic knockout for the mechanosensory proteins HOG1, WSC1, WSC2, WSC3, MYO1, MID2, TPM1, and TPM2. Yeast cells are cultured in Yeast Peptone Dextrose (YPD) medium (comprising 1% Yeast Extract (*w/v%*), 2% Peptone (*w/v%*) and 2% Dextrose (*w/v%*)) and cryopreserved as glycerol stocks (*30% v/v*). Prior to each experiment, we inoculate liquid YPD broth using frozen stocks at a 1% (*v/v*) seeding density and maintain the cells at 30° C for ∼20 h with continuous shaking to obtain a stationary growth phase culture. From these, we set a secondary culture using a 1% (*v/v*) inoculum in fresh YPD, which is incubated for ∼7 h at 30°C with continuous shaking to obtain an exponential phase culture. Cells from the exponential phase culture are used to inoculate 3D growth media for all assays, at a 5% (*v/v*) inoculum concentration, which is homogeneously dispersed throughout the 3D matrix. Further, for experiments involving mechanosensory mutants, all culture conditions are maintained as above with a singular exception – the inoculum volume from exponential phase secondary cultures to 3D matrix is normalized with respect to the wild-type in order to account for any inherent differences in the basal growth of mutant strains.

### 4.5. Modulation of osmotic pressure

For a limited set of experiments, we elevate the osmotic pressure of the liquid YPD broth to both 10-fold and 100-fold of the elastic modulus for the high confinement matrix. This is achieved by adding specific quantities of the bioinert polymer PEG-1000, as determined using the equation *π* = *icRT*, where *π* represents the osmotic pressure, *i* denotes the van’t Hoff index, *c* is the molar concentration of solute, R is the ideal gas constant, and T is the temperature in Kelvin.

### 4.6. Optical density measurements of growth in 3D matrices

Absorbance-based growth curve assays are performed using both liquid and 3D growth media inoculated with yeast cells as described above. For this, we acquire optical density measurements at a wavelength of 600nm using a multimode plate reader with built-in temperature control (Varioskan Lux). All values of absorbance are normalized by subtracting the initial time-point reading, which gives a measure of the net biomass increase with time.

### 4.7. Real-time imaging of yeast growth and cell health assessment

To visualize yeast growth under 3D confinement, we prepare samples consisting of homogeneously dispersed single cells as described above for the growth curve assays, which are housed in glass-bottom 35mm dishes and layered with oxygen-permeable mineral oil to prevent desiccation. Using a temperature-controlled microscope stage-mounted imaging chamber, we perform timelapse imaging of yeast growth for hundreds of cells in both low and high confinement matrices. For imaging cells in unconfined conditions, we aliquot a small volume of inoculated YPD broth onto a glass-bottom 35mm dish and sandwich the droplet by placing a YPD agar (1%) pad on top. To test whether confinement alters the cellular behavior, we retrieve cells embedded in the 3D matrix by diluting the 3D growth media with excess liquid YPD, vortexing to break up colonies, and using single cells obtained following this treatment to assess growth in unconfined condition (liquid YPD) as described above. We also assess the cell health by imaging fluorescently tagged mitochondria in cells maintained under unconfined, low, and high confinement matrices for 24 hours.

### 4.8. Colony and single-cell morphology analysis

To quantitatively analyze the colony morphology across both low and high confinement regimes, we acquire z-stack micrographs of cytoplasmic mCherry-expressing yeast colonies formed following 24 hours of growth under either condition. We use a custom MATLAB script to perform 3D segmentation of reconstructed colony images and quantify the 3D volume. Using the same samples, we obtain single cells by diluting the 3D growth media with liquid YPD and vortex mixing. A suspension containing these cells is spread flat on 1% YPD agar pads, flipped upside down to sandwich the cells between the pad and the base of a glass-bottom 35mm dish, and imaged. From these micrographs, we use built-in Fiji/ImageJ functions to characterize the area, perimeter, feret and circularity.

### 4.9. Validation of mechanosensory mutants

To verify the deletion of specific mechanosensory proteins, we first extract genomic DNA from cells grown in liquid YPD as previously described ^106^. Using primer pairs binding within the coding regions of these proteins as well as the housekeeping protein actin, we perform PCR amplifications to verify the presence or absence of mechanosensory protein-coding regions in these strains. Absence of amplification indicates a genomic knockout for the gene of interest, whereas presence of amplicons using the wild-type cells’ sample is used as a positive control. For the HOG1 knockout strain, we use a primer pair that amplifies a portion of the antibiotic cassette inserted in place of this gene’s coding region – hence in this case, a positive amplification indicates absence (i.e., knockout) of the gene of interest.

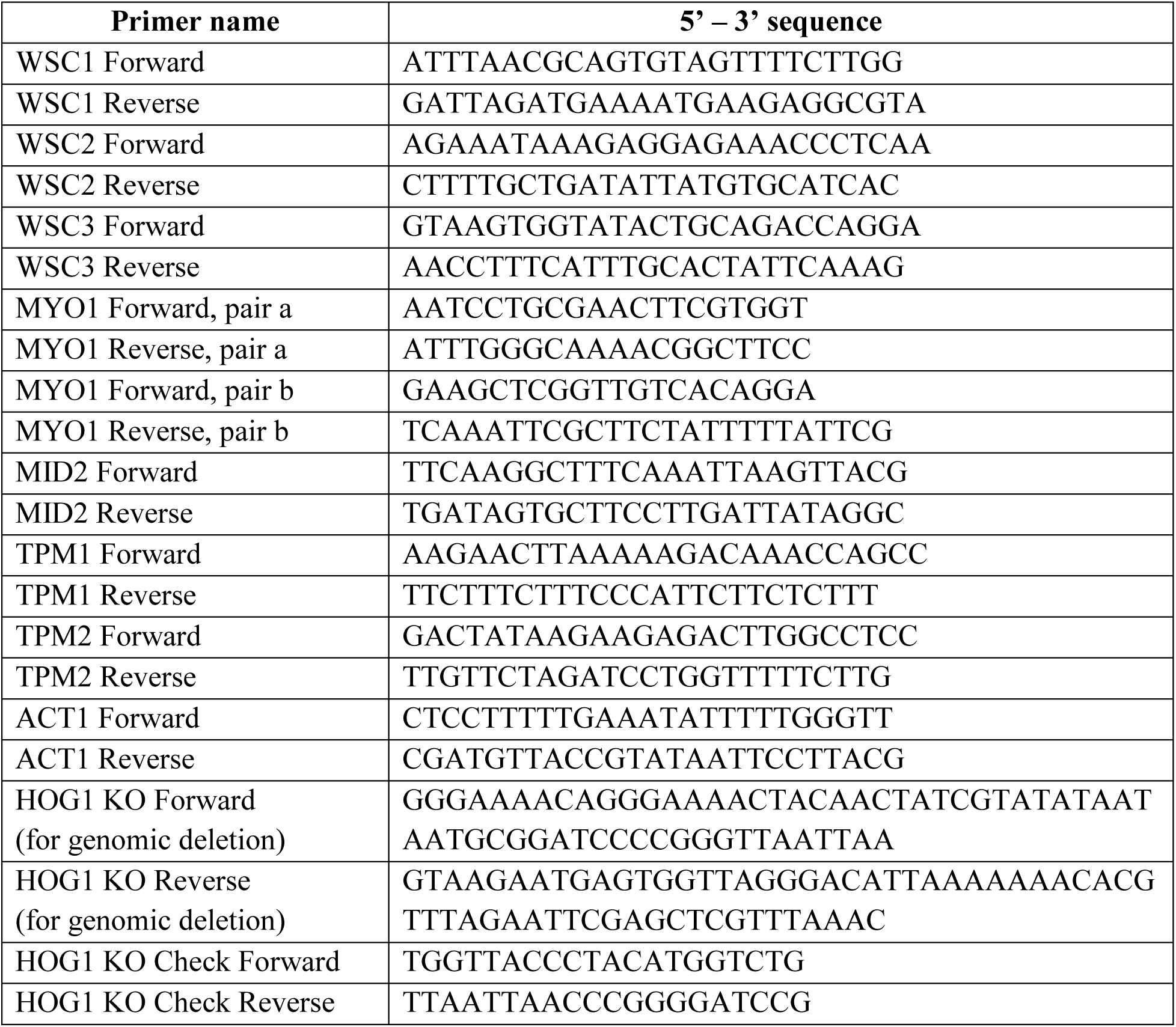

### 4.10. RNA extraction for bulk transcriptomics

We extract purified RNA from cells cultured for 24 hours within unconfined, low, and high confinement matrices using a hot acid phenol-based process. Briefly, a concentrated cell pellet is first obtained – for unconfined liquid samples, a direct centrifugation at 3000 xg for 1 minute suffices; whereas for cells embedded in 3D growth matrices, an initial dilution with excess liquid YPD, followed by repeated rounds of centrifugation at 3000 xg for 1 minute and subsequent washes with liquid YPD yields a concentrated cell pellet. We resuspend cell pellets obtained from 10 mL cultures in 400 μL of Tris-EDTA solution, to which 400 μL of hot acid phenol (pre-warmed to 65⁰C) is added. The mixture is vortexed for 10 minutes, and subsequently incubated at 65⁰C for 1 hour with intermittent vortexing for 1 minute every 5 minutes. Following a brief (5 minute) incubation on ice, we centrifuge the samples at 14000 xg for 10 minutes at 4⁰C. This results in a phase separation of the mixture into three distinct fractions – an upper aqueous phase, an interphase, and a lower organic phase. The RNA-containing upper aqueous fraction is carefully transferred to a fresh tube, wherein 400 μL of chloroform is added. This mixture is first vortexed for 3 minutes, followed by centrifugation at 14000 xg for 10 mins at 4⁰C. Once again, the upper aqueous fraction is retrieved, to which 400 μL of 3M sodium acetate and 1 mL of ice-cold 100% ethanol is added, and left to precipitate the RNA overnight at -20⁰C. Next day, following centrifugation at 14000 xg for 15 mins at 4⁰C, the supernatant is decanted, and the resultant pellet is washed twice with ice-cold 70% ethanol by first vortex mixing for 3 minutes and then centrifuging at 14000 xg for 15 mins at 4⁰C. Finally, the pellet is allowed to air-dry, and resuspended in 50 μL of RNase-free DEPC-treated water. Following quality assessments using Qubit Broad-Range RNA assays and fragment size analysis using Agilent Bioanalyzer, the samples are submitted for RNA-Seq library preparation.

### 4.11. RNA-Seq analysis

RNA samples are prepared using the NEBNext Poly(A) mRNA Magnetic Isolation Module (Catalog no-E7490L) and NEBNext® Ultra™ II Directional RNA Library Prep with Sample Purification Beads (Catalog no-E7765L). These samples are sequenced using an Illumina NovaSeq 6000 platform. Post sequencing, 23 to 36 million paired-end (2 * 101 bp) reads are obtained. FastQC v0.11.9 is used to perform the initial quality check. Adapters were trimmed from the reads using cutadapt v3.5 (-m 15 -j 0 -a AGATCGGAAGAGCACACGTCTGAACTCCAGTCA -A AGATCGGAAGAGCGTCGTGTAGGGAAAGAGTGT). The trimmed reads are mapped to the *Saccharomyces cerevisiae* genome (Saccharomyces_cerevisiae.R64-1-1 from Ensembl) using hisat2 v2.2.1. The mapping (sam) files are converted to bam using samtools 1.13. The mapped reads (95-98% mapping percentage) are counted using featureCounts v2.0.3. DESeq2 v1.34.0 is used to perform the read count normalisation and differential expression analysis. The plots are generated using R v4.1.0.2. Genes that show a fold change > 2, with an adjusted p-value <= 0.05, are considered as differentially expressed for further analysis. Gene ontology enrichment analysis of the differentially expressed genes is done using the R package ‘clusterProfiler v4.2.2’. Heatmaps were generated using the R package pheatmap, applying row-wise scaling to convert DESeq2-normalized counts into gene-wise Z-scores, which represent each gene’s expression relative to its mean, across samples for visualization.

### 4.12. Proteomics analysis

Cells are homogenized using bead beating in 100mM Tris.Cl buffer pH 8 and 5% SDS along with protease inhibitor cocktail. 25μg protein sample is first reduced with 5 mM TCEP and subsequently alkylated with 50 mM iodoacetamide. The protein is then digested using FASP with Trypsin at a 1:50 Trypsin-to-lysate ratio for 16 hours at 37°C. After digestion, the mixture is purified using a C18 silica cartridge and then concentrated by drying in a speed vac. The resulting dried pellet is resuspended in buffer A, which consists of 2% acetonitrile and 0.1% formic acid. For mass spectrometric analysis, all the experiments are performed on an Easy-nLC-1000 system (Thermo Fisher Scientific) coupled with an Orbitrap Exploris 240 mass spectrometer (Thermo Fisher Scientific) and equipped with a nano electrospray ion source. 1 μg of peptides’ sample dissolved in buffer A containing 2% acetonitrile/0.1% formic acid is resolved using Picofrit column (1.8-micron resin, 15cm length). Gradient elution is performed with a 0–38% gradient of buffer B (80% acetonitrile, 0.1% formic acid) at a flow rate of 500nl/min for 96mins, followed by 90% of buffer B for 11 min and finally column equilibration for 3 minutes. Orbitrap Exploris 240 is used to acquire MS spectra under the following conditions: Max IT = 60ms, AGC target = 300%; RF Lens = 70%; R = 60K, mass range = 375−1500. MS2 data are collected using the following conditions: Max IT= 60ms, R= 15K, AGC target 100%. MS/MS data are acquired using a data-dependent top20 method dynamically choosing the most abundant precursor ions from the survey scan, wherein dynamic exclusion is employed for 30s. Samples are processed and RAW files generated are analyzed with Proteome Discoverer (v2.5) against Uniprot reference database. For dual Sequest and Amanda search, the precursor and fragment mass tolerances are set at 10 ppm and 0.02 Da, respectively. The protease used to generate peptides, i.e. enzyme specificity is set for trypsin/P (cleavage at the C terminus of “K/R: unless followed by “P”). Carbamidomethyl on cysteine as fixed modification and oxidation of methionine and N-terminal acetylation are considered as variable modifications for database search. Both peptide spectrum match and protein false discovery rate are set to 0.01 FDR. For the comparison between cells grown in high confinement vs low confinement matrices, we set the statistical thresholds as adjusted p-value <= 0.05 and |log2FC| >= 0.5, with an additional quality control filter where only those proteins detected with high confidence across all replicates in both high confinement and low confinement samples are considered. We use a combination of Uniprot and SGD to identify annotated proteins amongst this filtered population, which represents the differentially abundant fraction of the proteome. The identified proteins are then subject to ontological classification using the SGD GO Slim Mapper for Biological Processes. A complete list of the annotated/identified differentially abundant proteins (totaling 112 unique proteins) is provided in Table S2. Z-score calculation for plotting the heatmap is performed using the definition of z-score as 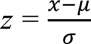, where *x* represents the abundance value of a single replicate from a given condition for each individual protein, while μ represents the overall mean and σ represents the overall standard deviation spanning all replicates for all proteins across both conditions.

### 4.13. Biophysical measurements of cell cycle phases

We quantitatively assess the inter-budding times, maturation times, puncta splitting times, puncta dispersion times, disassembly-assembly times, bud protrusion times, and bud size by manually tracking and segmenting cells in Fiji/ImageJ. For all calculations involving both wild-type and CDC10/CDC14-fusion reporter strains, we obtain a time difference between each iteration of the event from the time lapse imaging data. For measuring the bud sizes, we use in-built particle analysis functions in Fiji/ImageJ to obtain morphometric descriptors for the manual tracings. To calculate effective bud volume, we obtain the square root of the maximum projection of the bud area as an effective radius and use this to calculate the volume of an idealized sphere.

### 4.14. Statistical analyses

All statistical analyses shown in bar plots are performed using an unpaired t-test with a statistical threshold of p < 0.0001 for indicating significance. Sampling robustness is also evaluated for the primary phenotype of delayed interbudding times as well as maturation times. Here, we consider the entire sample space of either interbudding times or maturation times. From this, we randomly sample sub-populations, and measure the means, standard deviation, and standard error of such sub-populations. This process is iteratively repeated a hundred times in order to randomly sample a hundred such randomly chosen sub-populations from the overall dataset. Further, we perform this analysis for different sizes of sub-populations starting from n = 10 and increasing in increments of 10. These results are summarized in **SI Fig. 3.**

## Acknowledgments

We gratefully acknowledge Dr. Anjana Badrinarayanan for sharing the CDC14-mNeonGreen, CDC10-mScarlet, and mCherry reporter lines of *S. cerevisiae*. We thank Dr. Nivedita Hariharan, Insilytix Biosciences Pvt. Ltd. for assistance with the RNA-Seq analysis and graphical presentation. We are grateful to Dr. Saravanan Palani for sharing the SPC42-mNeonGreen, reporter line of *S. cerevisiae*. We acknowledge Prof. Sujit Datta for insightful discussions. We also acknowledge Aayushee Khanna, Ashitha B Arun, Meenakshi Ganesh, Lavanya Karinje, Sreesa Sreedharan, Swati Prasad, Shreyas Niphadkar, Vineeth Vengayil, and Akshaya Seshadri for their helpful advice and engagements in preliminary explorations associated with this work. We thank Dr. H. Krishnamurthy, Dr. Venkatesan Iyer, and the Central Imaging and Flow Cytometry Facility (CIFF) at NCBS for access to confocal microscopy. We thank Dr. Awadesh Pandit, Lakshminarayanan CP, and the Next Generation Sequencing Facility staff at NCBS for help with the RNA-Seq library preparation and sequencing runs. This work was supported by Nikon India.

## Author contributions

TB and MS conceptualized the study. MS with help from TB developed the methodology. MG, MS, and NB performed the experiments. MS, MG, and NB analyzed the data. MG, MS, and NB curated the data. NB and MS validated the key phenotypes. MS prepared the figures. MS with help from TB wrote the original manuscript. SL contributed critical inputs towards the study design and data interpretation, as well as provided feedback on the manuscript. SL shared critical reagents and strains for dissecting the cell cycle. TB secured funding and supervised the project.

## Data and materials availability

All data are available in the main text or the supplementary materials.

## Supplementary Materials for

**SI 1.**
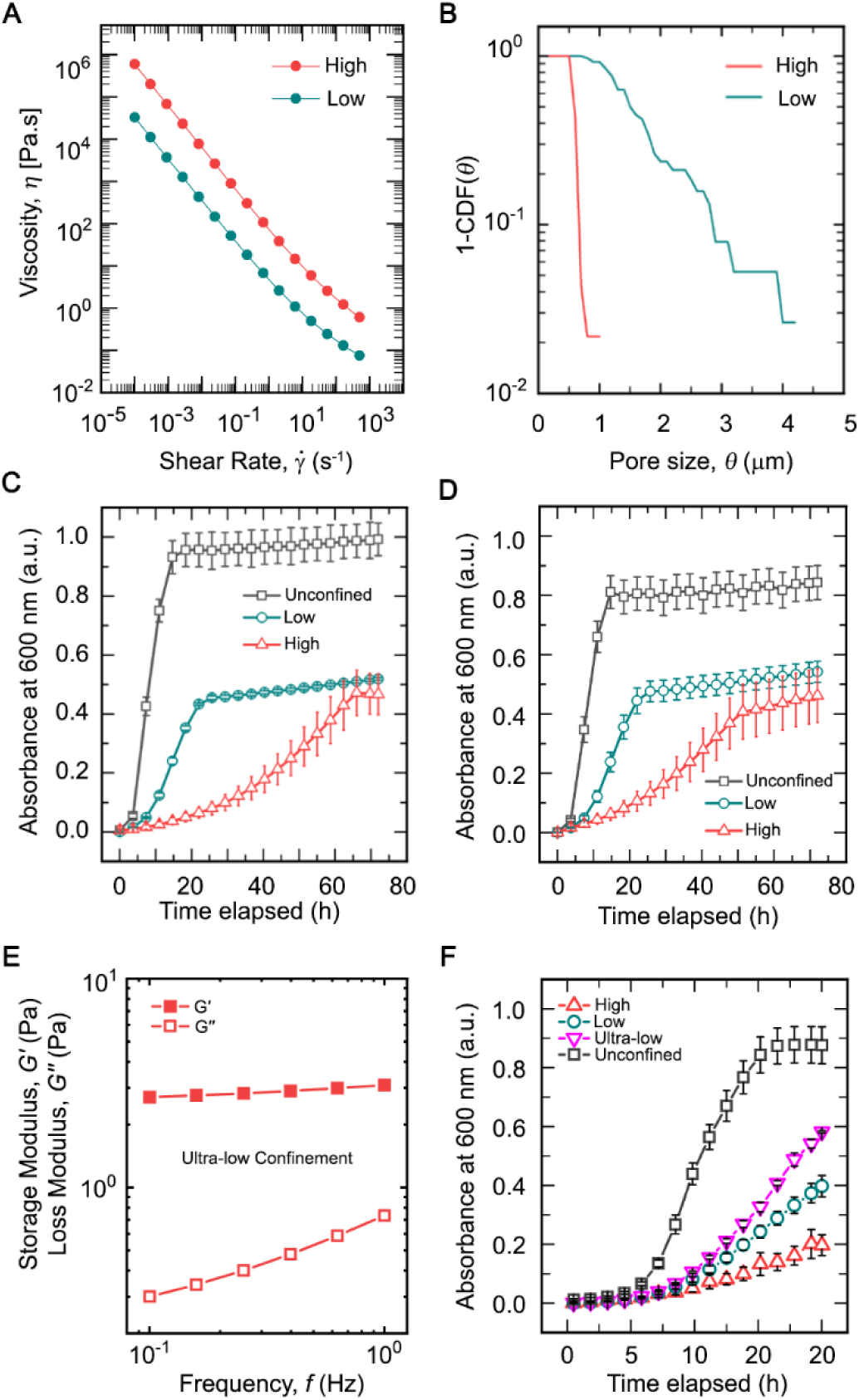
(**A**) Rheological measurements of the viscosity of low and high confinement 3D growth matrices, recorded across different shear rates. (**B**) Quantification of the 3D inter-particle pore spaces across low and high confinement 3D growth matrices, measured by tracking the diffusive motion of tracer particles and represented here as complementary cumulative distributions. (**C-D**) Independent biological replicates from optical density-based absorbance measurements of biomass production by budding yeast cells cultured under unconfined liquid as well as both low and high confinement 3D growth media. mean +/- s.d., n = 3 technical replicates for each condition. (**E**) Oscillatory shear rheology measurements performed on ultra-low confinement matrices to evaluate the material’s shear moduli reveal the elastic storage modulus as being significantly greater than the viscous loss modulus, suggesting that the 3D growth media exhibits soft solid-like properties. (**F**) Optical density-based absorbance measurements of biomass production by budding yeast cells cultured under ultra-low confinement, contrasted against data from unconfined liquid as well as both low and high confinement 3D growth media. mean +/- s.d., n = 3 technical replicates for each condition.

**SI 2.**
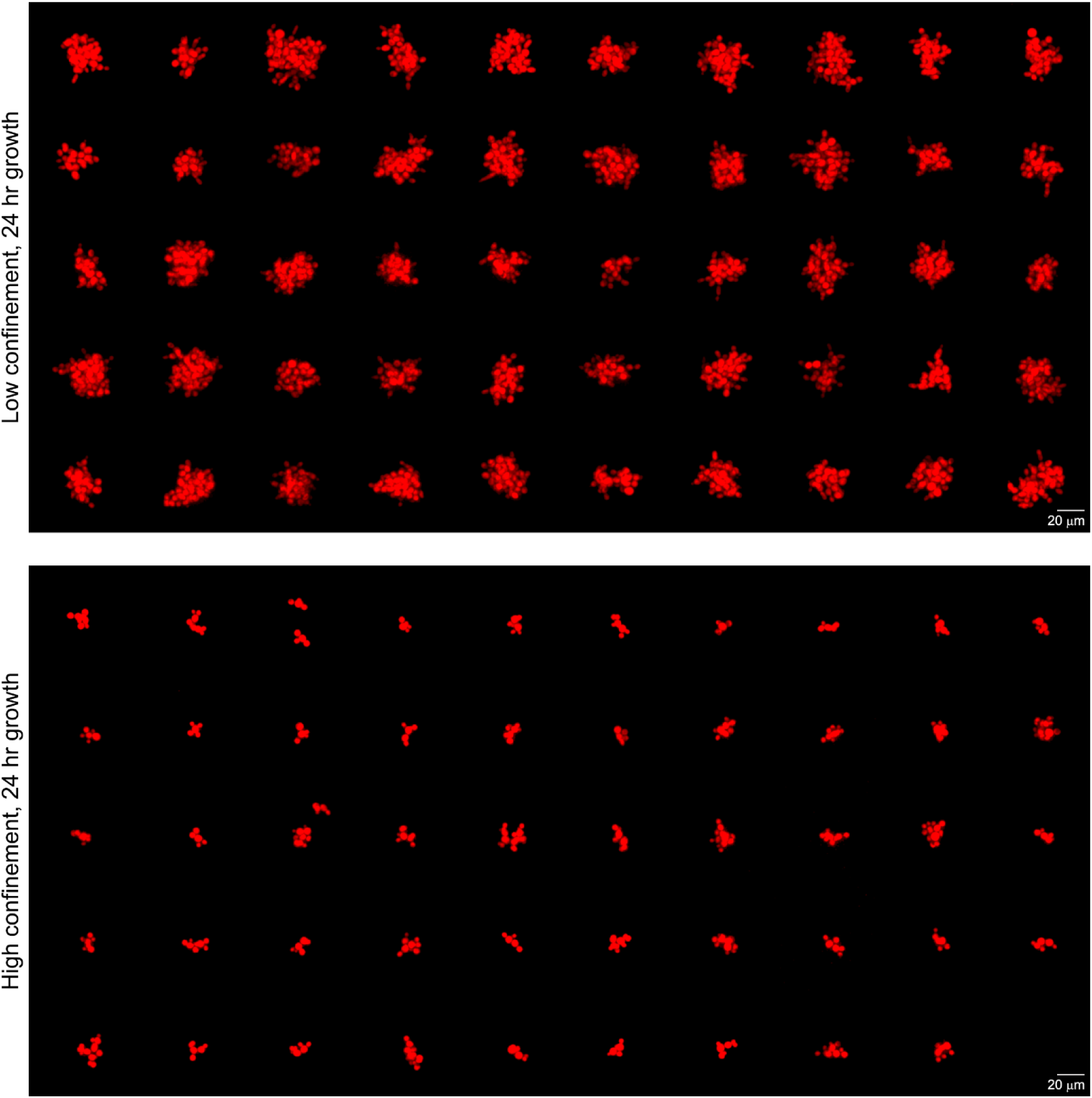
Representative micrographs showcasing 3D colonies of cytoplasmic mCherry-expressing budding yeast, cultured under either low or high degrees of physical confinement for 24 hours within the 3D growth media.

**SI 3.**
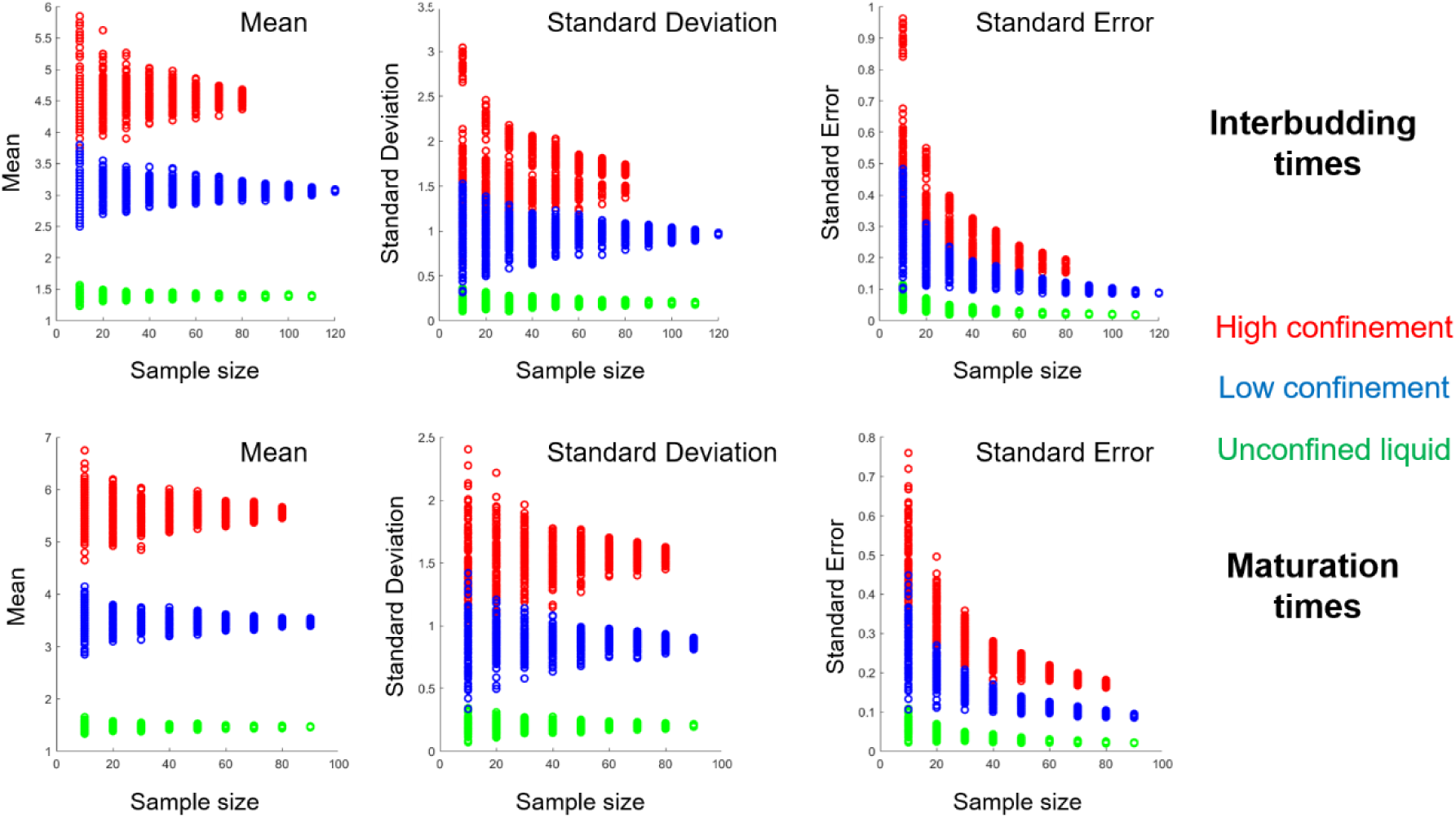
Statistical analyses to evaluate the population sampling robustness by randomized and iterative sub-sampling of the interbudding and maturation times’ datasets, followed by assessing the measures of central tendency and variance.

**SI 4.**
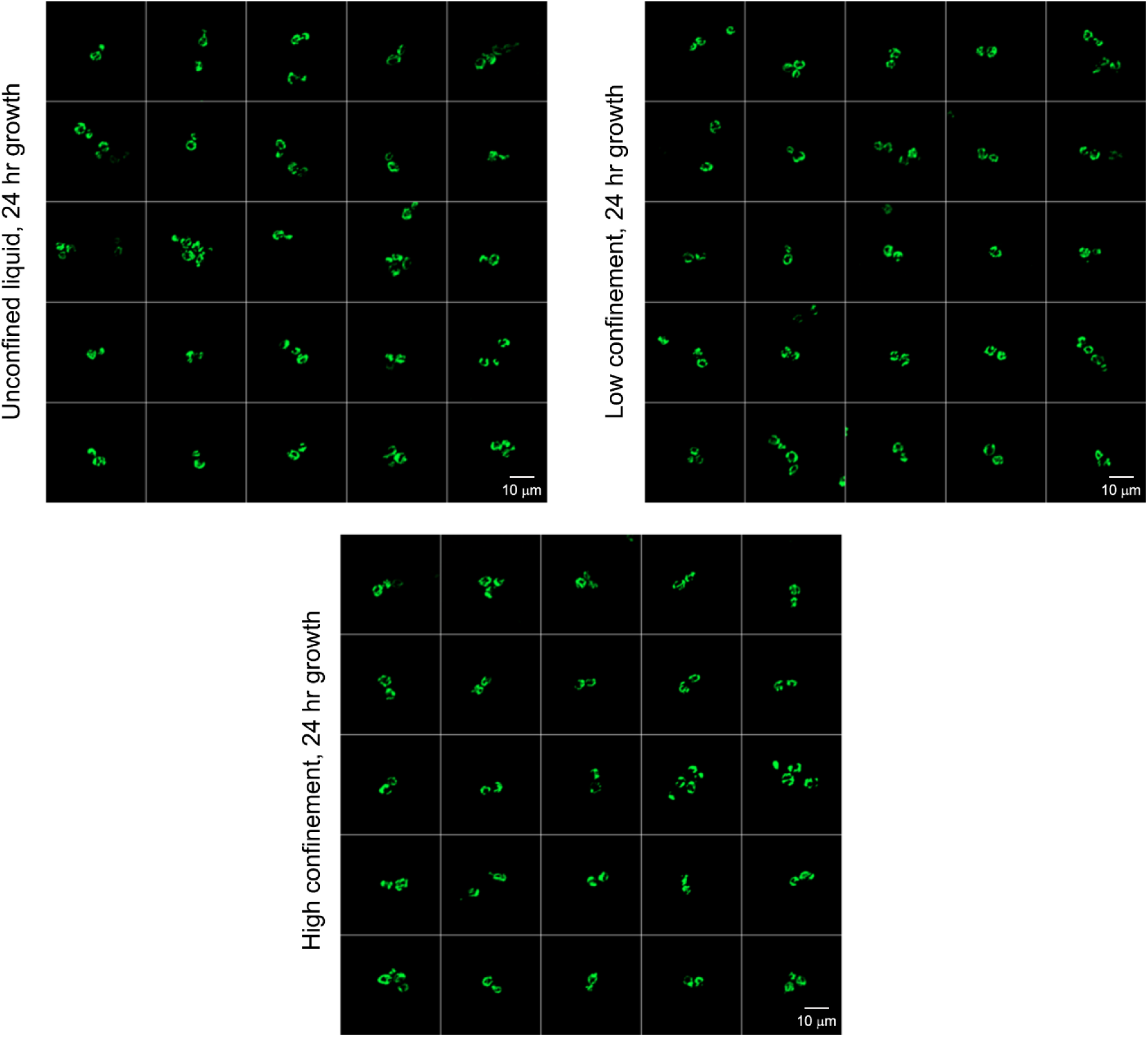
Growth under either unconfined liquid, low confinement, or high confinement does not appreciably alter the mitochondrial morphology, suggesting that prolonged budding times under higher confinement does not differentially disrupt the cell metabolism.

**SI 5.**
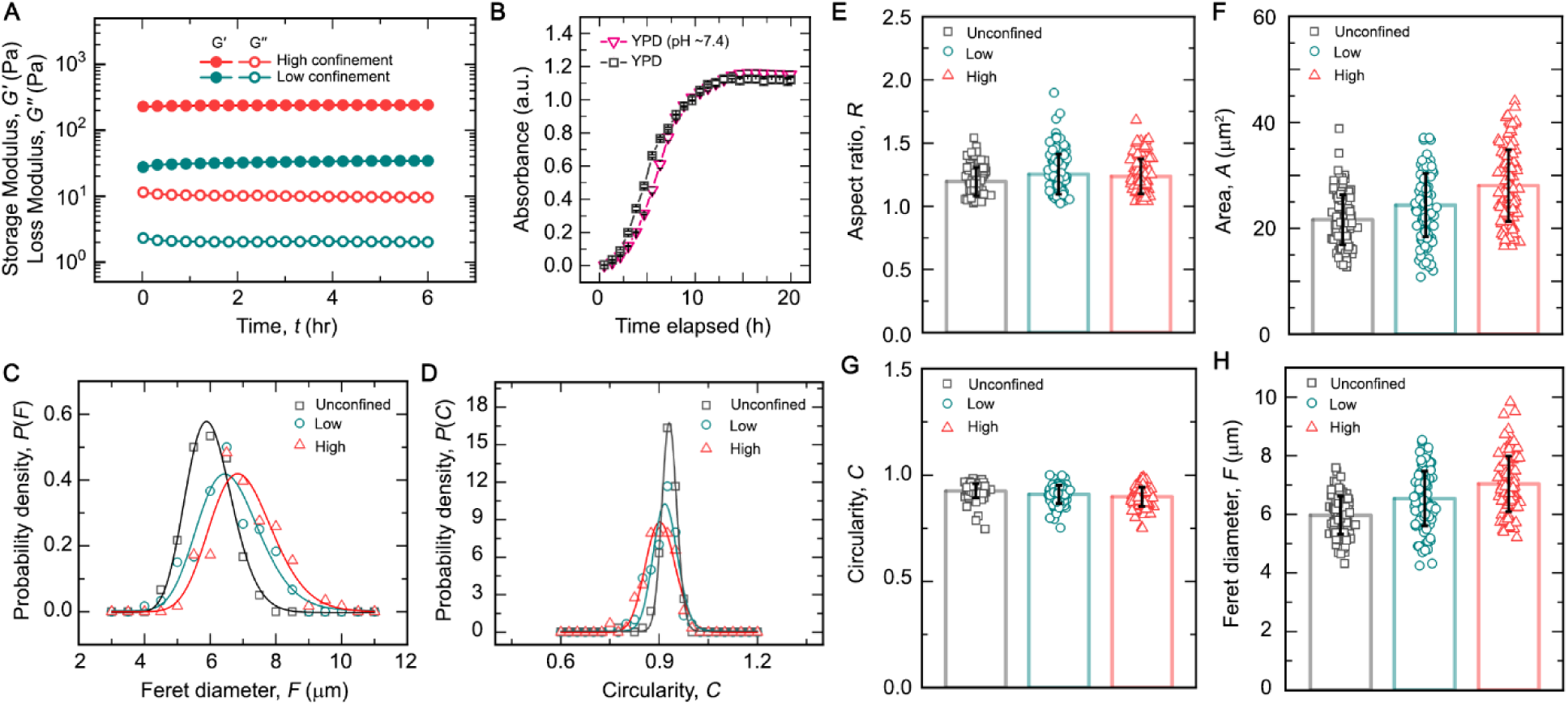
(**A**) Continuous measurement of the rheological properties of the 3D growth media over 6 hours of growth at 30°C. (**B**) Shaken liquid cultures assaying the growth of yeast cells in standard YPD (pH ∼6.5) vs YPD with an initially-adjusted pH of ∼7.4. mean +/- s.d., n = 3 technical replicates for each condition. (**C** and **D**) Probability density distributions from single-cell morphometrical measurements of feret diameter (the largest dimension spanning the cell body) and circularity obtained from cells cultured for 24 hours under unconfined liquid, as well as both low and high degrees of 3D confinement. (**E-H**) Single-cell measurements detailing various morphometric descriptors - aspect ratio, area, circularity, and feret diameter - of cells cultured for 24 hours under unconfined liquid as well as both low and high degrees of 3D confinement. mean +/- s.d., n >= 100 individual cells for each condition.

**SI 6.**
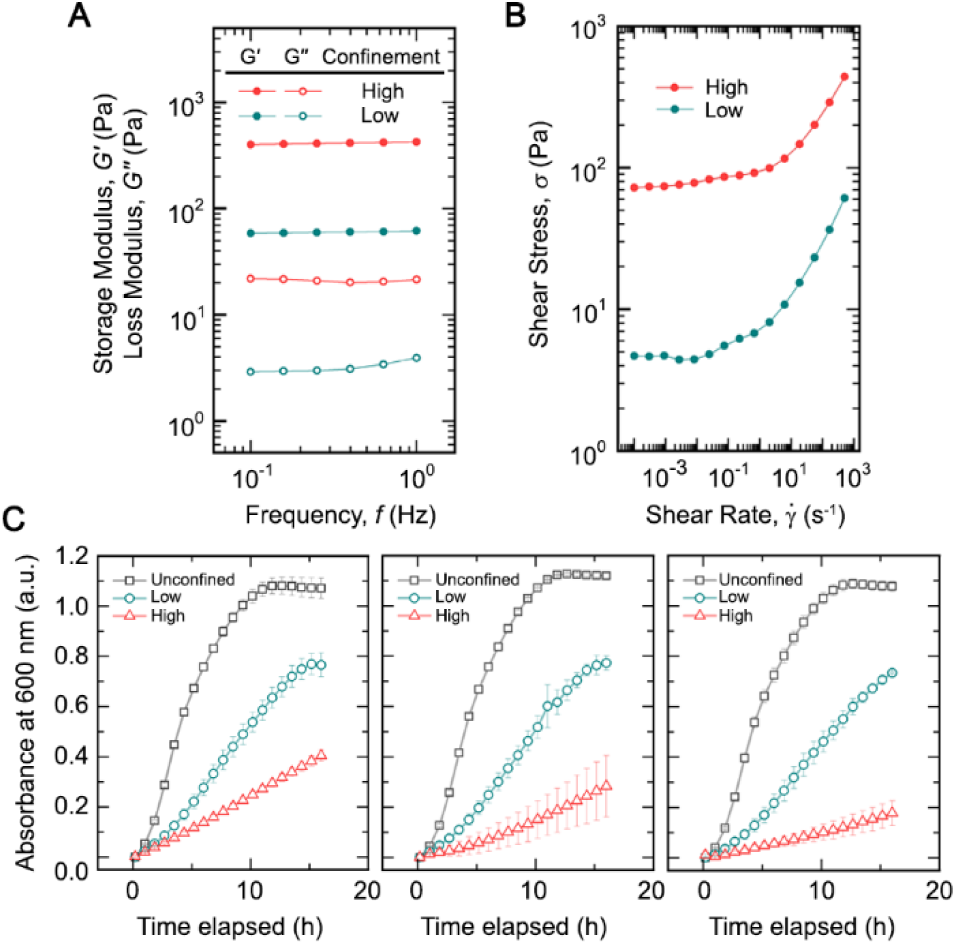
Rheological measurements of (**A**) shear moduli - elastic storage modulus and viscous loss modulus - as well as (**B**) the shear response to varying shear rates for both low and high confinement ETD2020-based 3D matrices. (**C**) Optical density-based absorbance measurements of biomass production by budding yeast cells cultured under unconfined liquid, as well as both low and high degrees of confinement within the ETD2020-based 3D growth media. mean +/- s.d., n = 3 technical replicates for each condition.

**SI 7.**
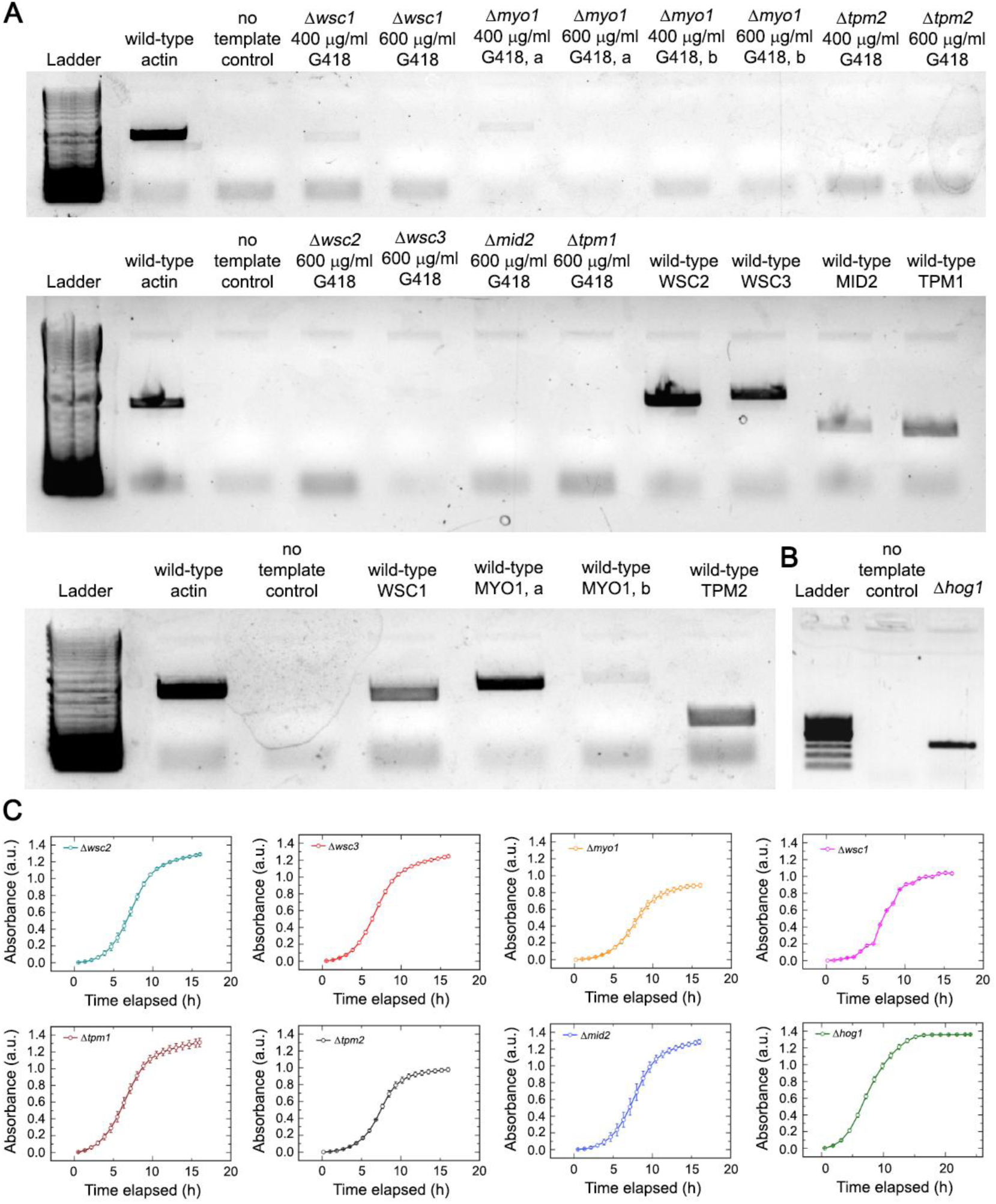
(**A**) Electrophoresis gels showing PCR amplicons for the following genomic locii - WSC1, MYO1, TPM2, WSC2, WSC3, MID2, and TPM1. Template genomic DNA sources are either from specific genomic knockout strains or the wild-type strain. Actin from wild-type strain is shown as a positive control. Two degrees of G418 antibiotic selection were applied on some of the genomic knockout strains, and for all experiments, we used populations selected under the higher 600 μg/ml antibiotic pressure. For MYO1, we employ two different primer pairs, denoted by a and b respectively. (**B**) Electrophoresis gel showing a PCR amplicon for the antibiotic resistance cassette inserted in place of the region coding for HOG1 – indicative of a genomic knockout. (**C**) Shaken liquid cultures assaying the baseline growth behavior of the eight knockout strains. mean +/- s.d., n = 3 technical replicates for each condition.

**SI 8.**
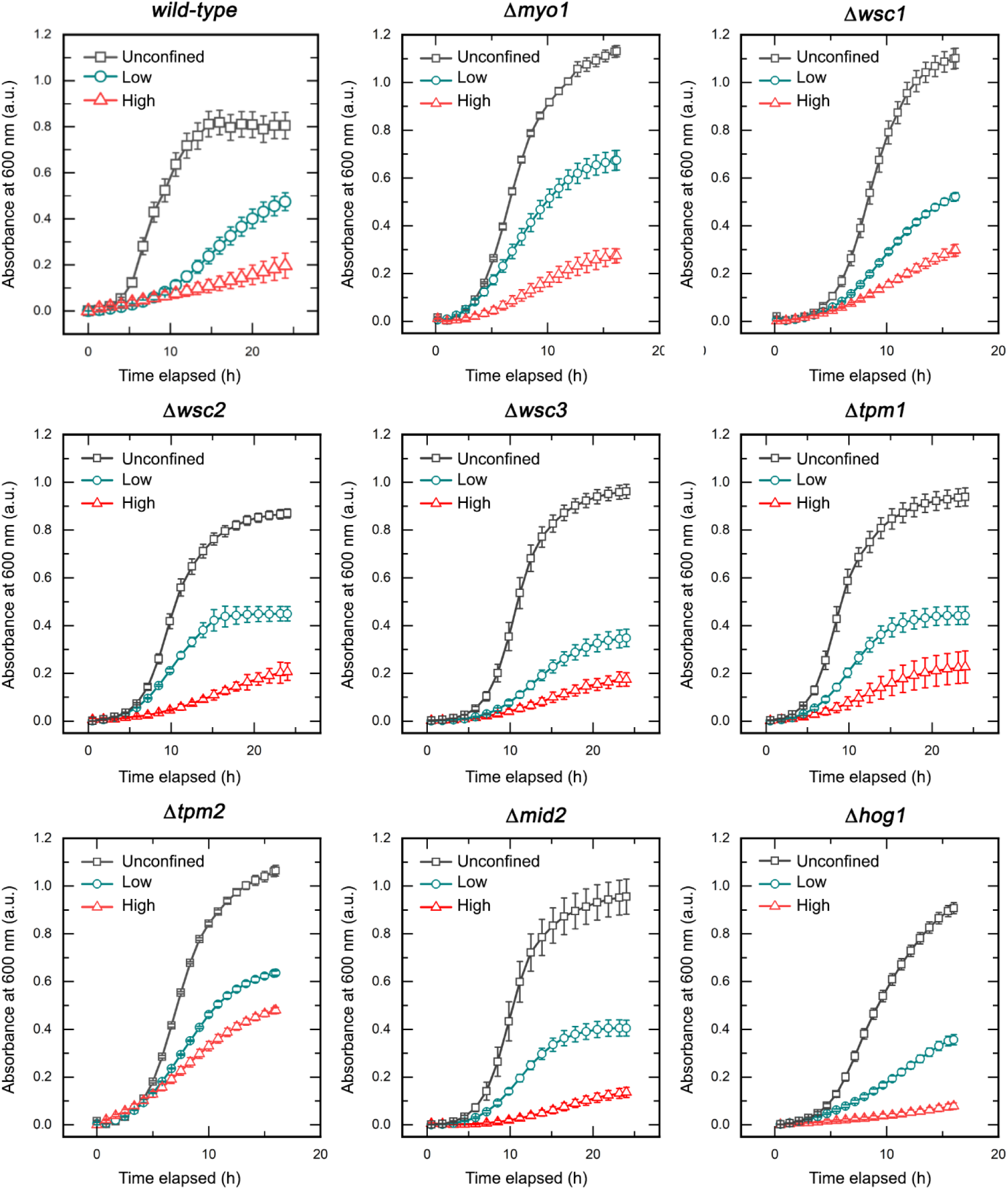
Growth curve assays performed across different degrees of confinement for the wild-type strain and the eight mechanosensory mutant strains.

**SI 9.**
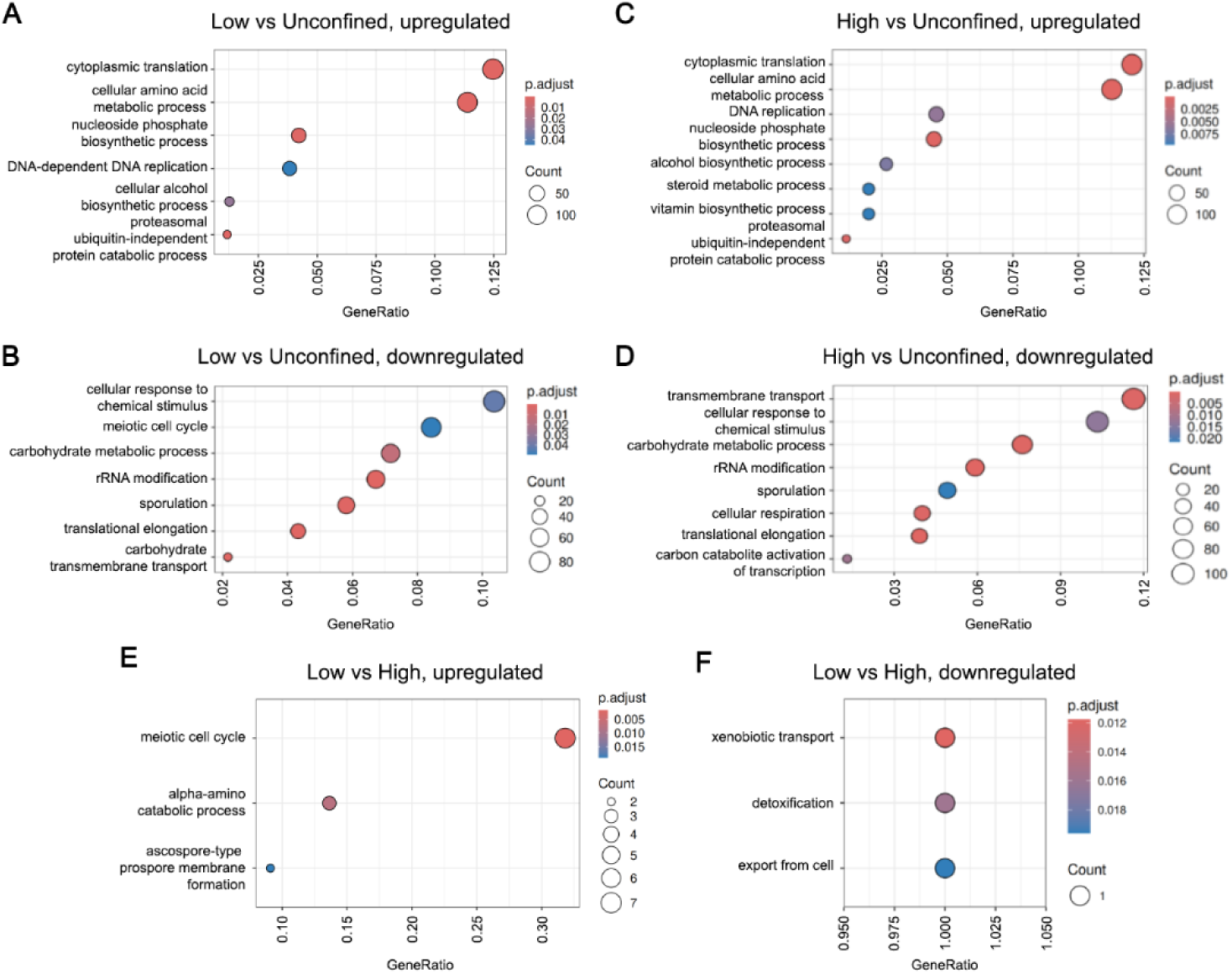
Gene ontology dot plots for Biological Processes classification, obtained from pairwise differential expression analyses between (**A, C,** and **E**) upregulated groups across low confinement vs unconfined liquid, high confinement vs unconfined liquid, and low vs high confinement, as well as (**B, D,** and **F**) downregulated groups across low confinement vs unconfined liquid, high confinement vs unconfined liquid, and low vs high confinement.

**SI 10.**
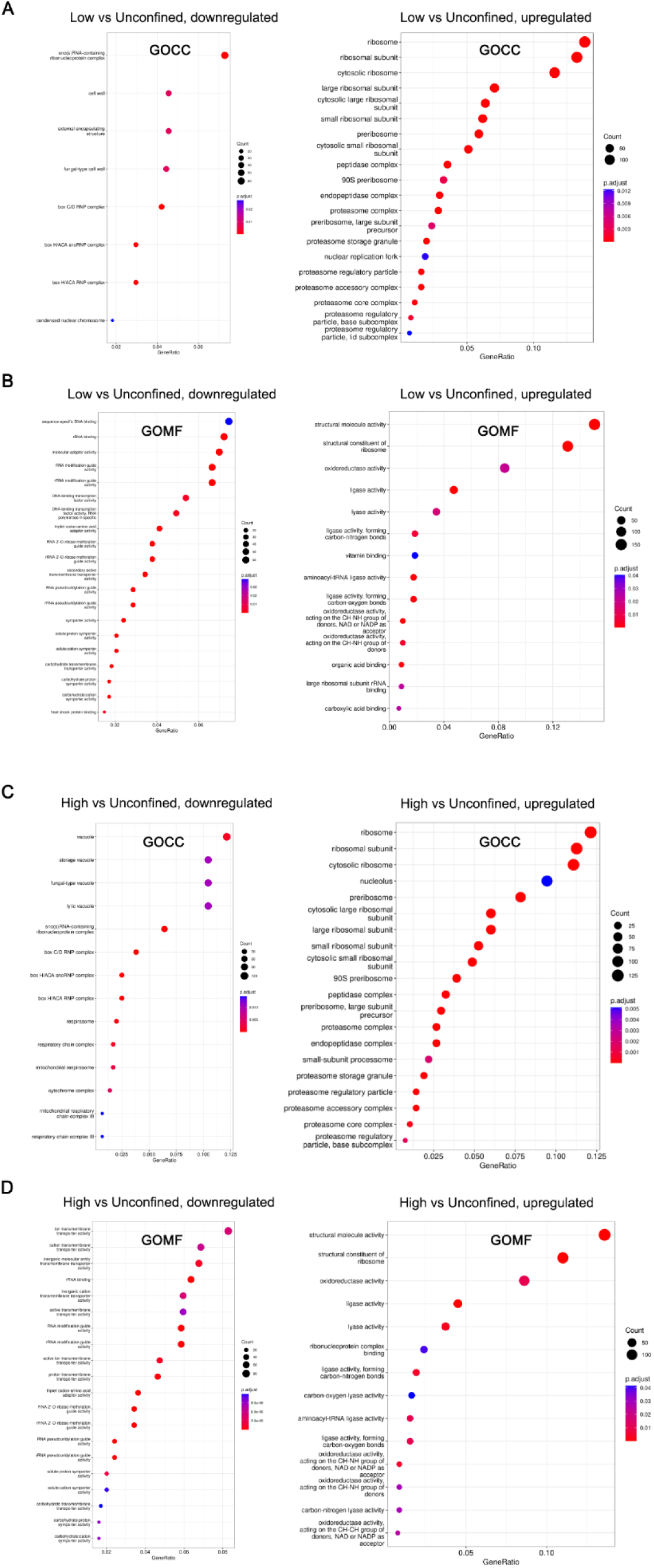
Dot plots showing downregulated and upregulated groups across (**A** and **C**) Gene Ontology Cellular Components (GOCC) and (**B** and **D**) Gene Ontology Molecular Functions (GOMF) classifications, obtained from pairwise differential expression analyses between cells grown in either (**A** and **B**) low confinement vs unconfined liquid, or (**C** and **D**) high confinement vs unconfined liquid.

**SI 11.**
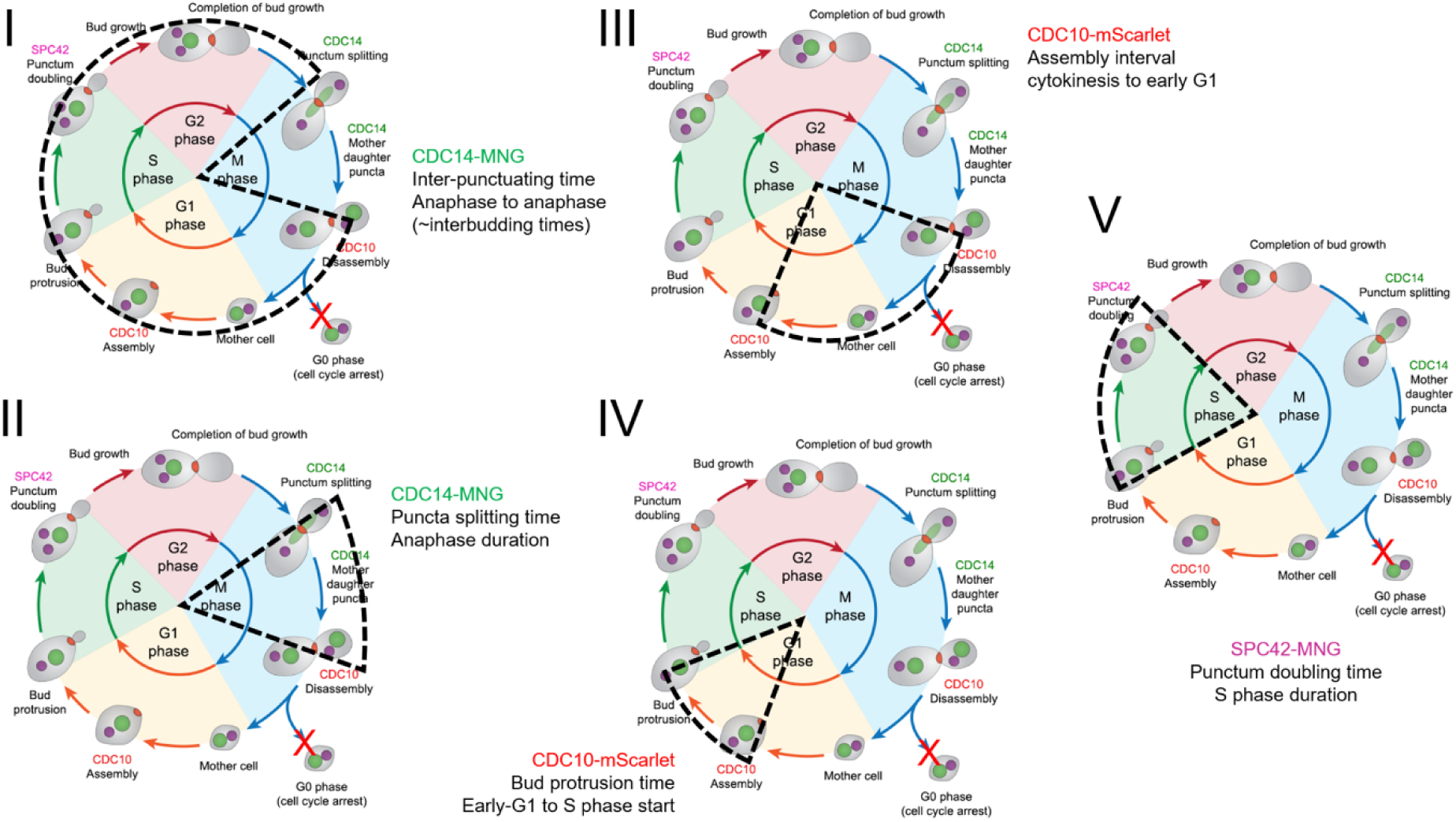
Schematic illustration of cell cycle dissection using fluorescently-tagged reporter constructs.

**SI 12.**
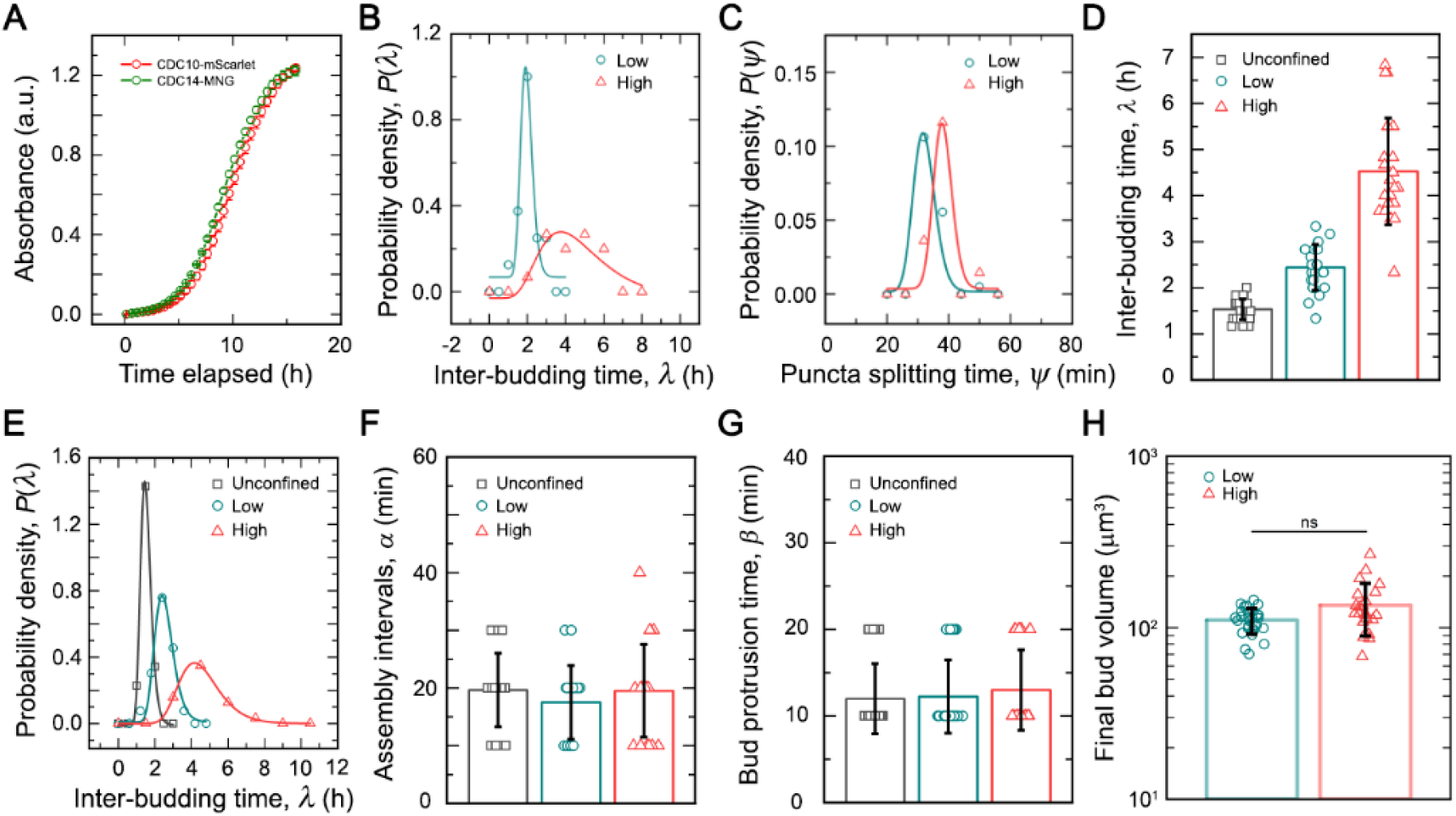
(**A**) Optical density-based absorbance measurements performed on shaken liquid cultures of CDC14-mNeonGreen and CDC10-mScarlet for assaying the baseline growth dynamics of fluorescent reporter strains. Mean +/- s.d., n = 3 technical replicates for each condition. (**B** and **C**) Probability density distributions obtained from measurements of inter-budding times and puncta splitting times from time lapse imagings of the CDC14-mNeonGreen reporter strain cultured under both low and high degrees of 3D confinement. n >= 15 individual events for each condition. (**D** and **E**) Inter-budding times, mean +/- s.d., n >= 20 individual events for each condition. (**F**) disassembly-assembly interval measurements, mean +/- s.d., n >= 20 individual events for each condition, and (**G**) bud protrusion times obtained from timelapse imaging of CDC10-mScarlet cultured under unconfined liquid, as well as both low and high degrees of 3D confinement. mean +/- s.d., n >= 40 individual events for each condition. (**H**) Quantification of the final bud volumes across different mechanical regimes. mean +/- s.d., n >= 22 individual cells for each condition. Statistical significance calculated using an unpaired t test (ns indicates not significant, p > 0.01).

**SI 13.**
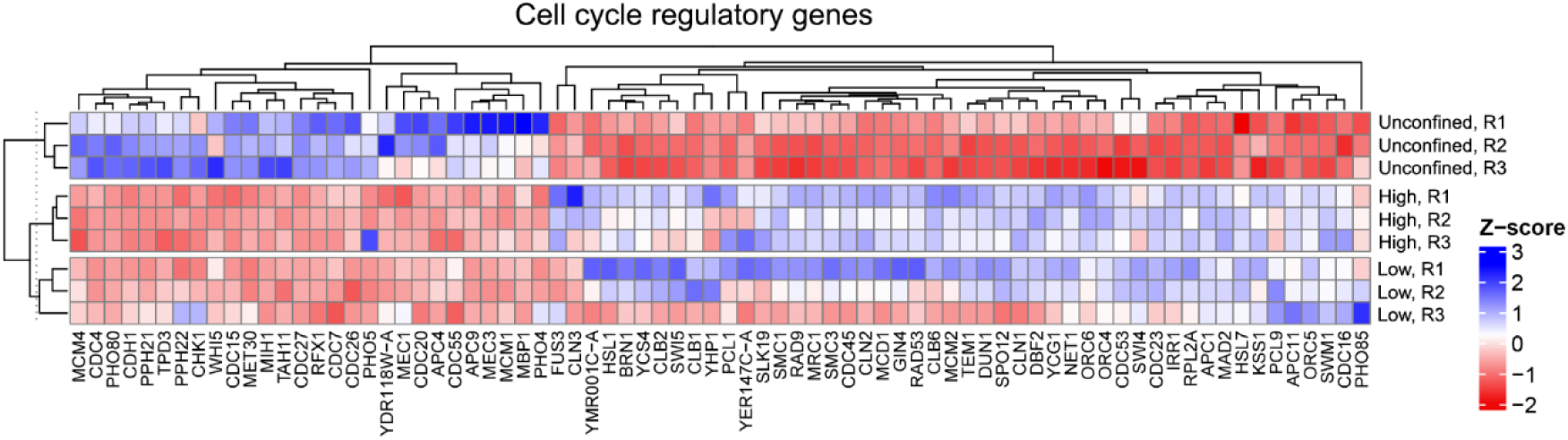
Heatmap detailing relative expression levels for genes involved in cell cycle regulation.

**SI 14.**
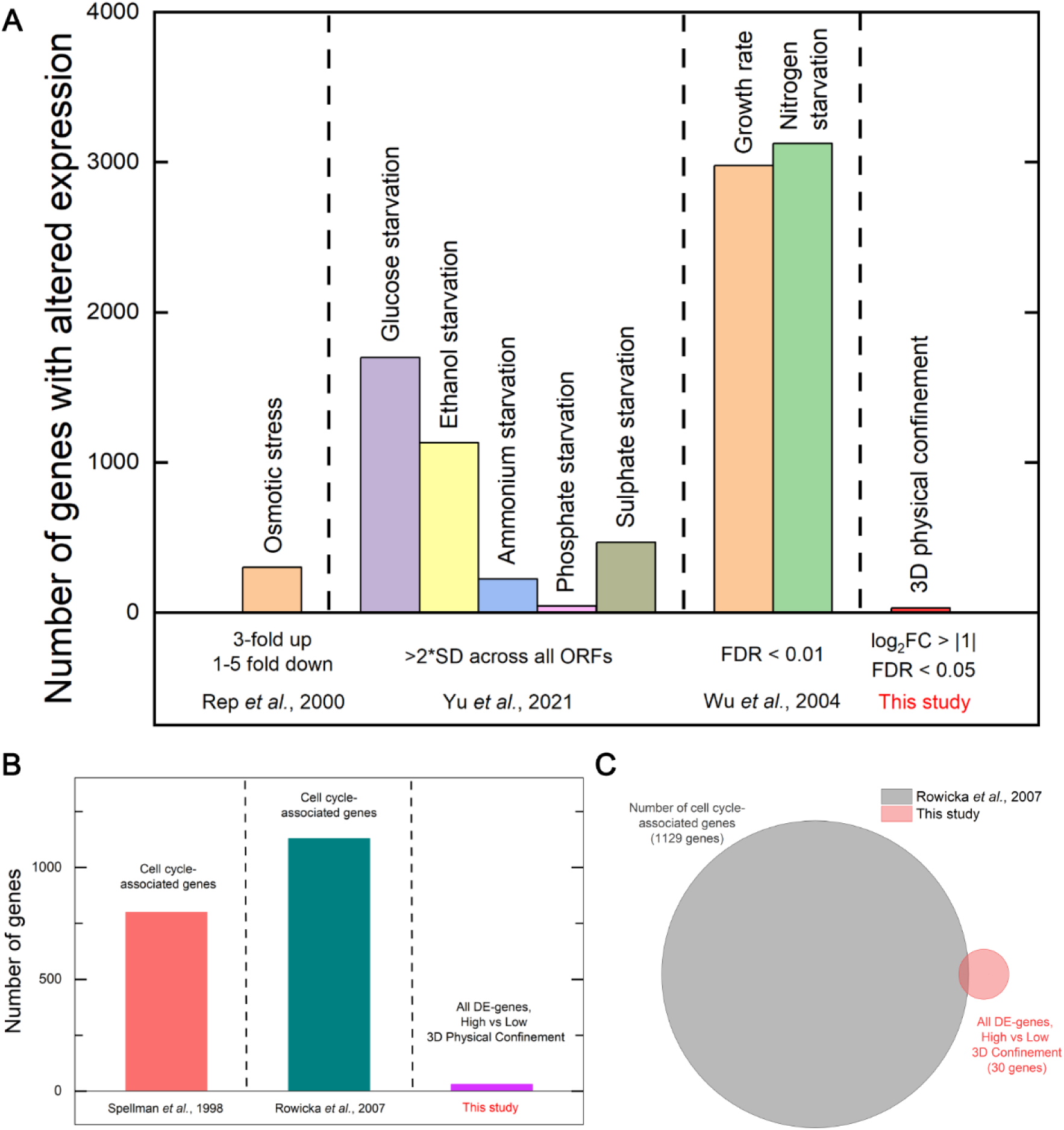
(**A**) Comparative analysis of RNA-level expression changes reported for budding yeast cells exposed to different types of perturbations^1–3^, which are known to also arrest/delay cell cycle progression. (**B** and **C**) Prior analyses of genes associated with cell cycle regulation^4,5^ compared against all the genes differentially expressed across cells grown under high confinement matrices vs low confinement matrices. The minimal overlap between these clearly demonstrates that cell cycle regulation is not specifically perturbed under high confinement.

**Table S1.**
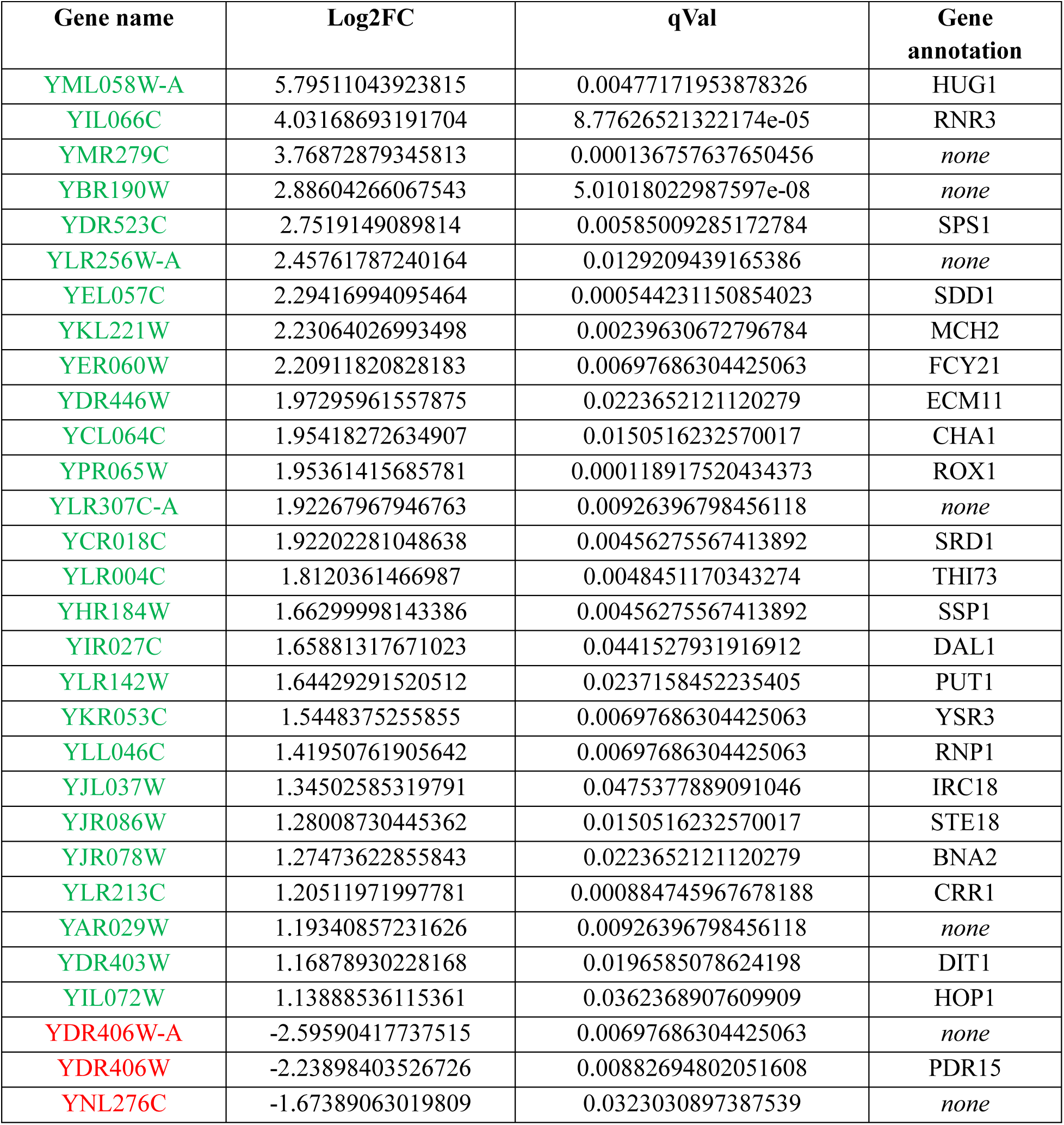
List of differentially regulated genes compared between cells grown under low confinement vs high confinement 3D growth matrices, subject to the condition that |log2FC| >= 1 and qVal (adjusted p-value) <= 0.05. Green text indicates upregulated genes, whereas red text indicates downregulated genes.

**Table S2.** List of annotated genes identified based on differentially abundant protein expression profiles, compared between cells grown under high confinement vs low confinement 3D growth matrices, subject to the condition that |log2FC| >= 0.5 and adjusted p-value <= 0.05. Green text indicates upregulated genes, whereas red text indicates downregulated proteins.

| SGD ID | Log2FC | Adjusted p-value | SGD Gene |
| --- | --- | --- | --- |
| YIL021W | -6.93448 | 0.007344 | RPB3 |
| YLR275W | -5.3287 | 0.002727 | SMD2 |
| YKR025W | -4.57832 | 0.005496 | RPC37 |
| YHR107C | -2.67617 | 0.00541 | CDC12 |
| YJL096W | -2.31658 | 0.010572 | MRPL49 |
| YDR101C | -2.12231 | 0.007742 | ARX1 |
| YBR072W | -2.09908 | 0.002727 | HSP26 |
| YDL204W | -2.00975 | 0.034625 | RTN2 |
| YMR276W | -1.98531 | 0.007742 | DSK2 |
| YLR120C | -1.96668 | 0.015602 | YPS1 |
| YOR360C | -1.91214 | 0.00334 | PDE2 |
| YLR175W | -1.90836 | 0.005496 | CBF5 |
| YLR186W | -1.7752 | 0.019208 | EMG1 |
| YDL084W | -1.72849 | 0.013632 | SUB2 |
| YGL171W | -1.65954 | 0.040597 | ROK1 |
| YMR256C | -1.58342 | 0.014144 | COX7 |
| YJL052W | -1.362 | 0.013666 | TDH1 |
| YER103W | -1.33761 | 0.015628 | SSA4 |
| YDR117C | -1.2666 | 0.013632 | TMA64 |
| YJL033W | -1.26591 | 0.02483 | HCA4 |
| YNL160W | -1.25954 | 0.036361 | YGP1 |
| YHR039C | -1.24424 | 0.045084 | MSC7 |
| YBL015W | -1.22489 | 0.047159 | ACH1 |
| YEL050C | -1.21105 | 0.035799 | RML2 |
| Q0250 | -1.20558 | 0.039424 | COX2 |
| YGL012W | -1.1991 | 0.019085 | ERG4 |
| YML028W | -1.17545 | 0.028803 | TSA1 |
| YML128C | -1.16532 | 0.012554 | MSC1 |
| YGL068W | -1.14661 | 0.028297 | MNP1 |
| YIL115C | -1.07633 | 0.013632 | NUP159 |
| YER014W | -1.06979 | 0.035252 | HEM14 |
| YHR138C | -1.04743 | 0.049138 | *none* |
| YNL015W | -1.038 | 0.007742 | PBI2 |
| YDR529C | -1.02952 | 0.013666 | QCR7 |
| YPR139C | -1.00444 | 0.01966 | LOA1 |
| YDR204W | -0.99654 | 0.013666 | COQ4 |
| YBR299W | -0.95993 | 0.02016 | MAL32 |
| YLR106C | -0.95748 | 0.035501 | REA1 |
| YGL231C | -0.94819 | 0.027126 | EMC4 |
| YHL021C | -0.93376 | 0.028297 | AIM17 |
| YHR087W | -0.89608 | 0.015678 | RTC3 |
| YNL200C | -0.89025 | 0.024475 | NNR1 |
| YDR343C | -0.88885 | 0.013666 | HXT6 |
| YFR053C | -0.84299 | 0.031291 | HXK1 |
| YGR245C | -0.79672 | 0.023239 | SDA1 |
| YPR160W | -0.78485 | 0.017642 | GPH1 |
| YJL183W | -0.77037 | 0.048531 | MNN11 |
| YGR149W | -0.72471 | 0.048531 | GPC1 |
| YFL030W | -0.71391 | 0.044952 | AGX1 |
| YJL164C | -0.70736 | 0.033899 | TPK1 |
| YHR104W | -0.68894 | 0.028297 | GRE3 |
| YNR001C | -0.66477 | 0.033899 | CIT1 |
| YLL036C | -0.66057 | 0.049138 | PRP19 |
| YKL122C | -0.65886 | 0.049138 | SRP21 |
| YDR353W | -0.64638 | 0.039061 | TRR1 |
| YNL081C | -0.64189 | 0.039424 | SWS2 |
| YHR001W-A | -0.63913 | 0.048531 | QCR10 |
| YOL058W | 0.573726 | 0.049138 | ARG1 |
| YPR035W | 0.575151 | 0.038727 | GLN1 |
| YBL017C | 0.580531 | 0.042969 | PEP1 |
| YDR341C | 0.585156 | 0.045013 | *none* |
| YKL211C | 0.607197 | 0.041785 | TRP3 |
| YNL104C | 0.61513 | 0.027712 | LEU4 |
| YBR115C | 0.624269 | 0.048531 | LYS2 |
| YIR034C | 0.634508 | 0.030773 | LYS1 |
| YLR056W | 0.64451 | 0.028297 | ERG3 |
| YMR134W | 0.646072 | 0.03918 | ERG29 |
| YHR018C | 0.664172 | 0.035463 | ARG4 |
| YBR096W | 0.704685 | 0.041785 | *none* |
| YBR249C | 0.721427 | 0.034625 | ARO4 |
| YOL090W | 0.742702 | 0.040339 | MSH2 |
| YJL088W | 0.749306 | 0.040339 | ARG3 |
| YBR104W | 0.755165 | 0.034625 | YMC2 |
| YBR177C | 0.778602 | 0.041785 | EHT1 |
| YDL182W | 0.814501 | 0.019085 | LYS20 |
| YNR050C | 0.83053 | 0.047159 | LYS9 |
| YMR090W | 0.845051 | 0.019085 | *none* |
| YDR481C | 0.861924 | 0.012353 | PHO8 |
| YKL120W | 0.892747 | 0.042583 | OAC1 |
| YJR130C | 0.910169 | 0.015219 | STR2 |
| YDR427W | 0.924682 | 0.01966 | RPN9 |
| YLL028W | 0.943025 | 0.040339 | TPO1 |
| YRB30 | 0.973658 | 0.015678 | YRB30 |
| YBR176W | 0.976399 | 0.036361 | ECM31 |
| YGR207C | 0.991902 | 0.015602 | CIR1 |
| YOR069W | 0.992361 | 0.012554 | VPS5 |
| YLR083C | 1.081498 | 0.015602 | EMP70 |
| YDL131W | 1.113874 | 0.027479 | LYS21 |
| YNL323W | 1.135942 | 0.03673 | LEM3 |
| YER175C | 1.14617 | 0.012353 | TMT1 |
| YMR310C | 1.147512 | 0.048531 | UPA2 |
| YER090W | 1.196462 | 0.013666 | TRP2 |
| YHR137W | 1.248256 | 0.010095 | ARO9 |
| YEL022W | 1.265405 | 0.034625 | GEA2 |
| YDR007W | 1.268959 | 0.006998 | TRP1 |
| YDR310C | 1.324872 | 0.015911 | SUM1 |
| YKL033W-A | 1.358508 | 0.022818 | *none* |
| YMR095C | 1.379381 | 0.00541 | SNO1 |
| YDL085W | 1.419685 | 0.048531 | NDE2 |
| YEL046C | 1.528509 | 0.038291 | GLY1 |
| YPL058C | 1.591253 | 0.010572 | PDR12 |
| YOR215C | 1.59293 | 0.015602 | AIM41 |
| YHR052W | 1.594602 | 0.008643 | CIC1 |
| YDR011W | 1.695826 | 0.002727 | SNQ2 |
| YPL118W | 1.713338 | 0.007344 | MRP51 |
| YNR069C | 1.791853 | 0.012353 | BSC5 |
| YDR345C | 1.862597 | 0.002727 | HXT3 |
| YDR175C | 1.975932 | 0.002727 | RSM24 |
| YIL030C | 1.97611 | 0.00541 | SSM4 |
| YLR051C | 2.107179 | 0.048531 | FCF2 |
| YHR216W | 2.349408 | 0.048352 | IMD2 |
| YGR220C | 2.906551 | 0.002523 | MRPL9 |

**Table S3.** Ontological classifications of biological processes being either downregulated or upregulated between cells grown under high confinement vs low confinement 3D matrices, represented as the absolute number of differentially abundant proteins identified as belonging to each ontological grouping compared against the total number of members in each such grouping.

| Upregulated Biological Processes |  | Downregulated Biological Processes |  |
| --- | --- | --- | --- |
| Ontological Term | No. of proteins | Ontological Term | No. of proteins |
| amino acid metabolic process | 16 out of 161 total entries | cellular respiration | 6 out of 112 total entries |
| transmembrane transport | 5 out of 277 total entries | rRNA processing | 6 out of 335 total entries |
| intracellular monoatomic ion homeostasis | 3 out of 132 total entries | carbohydrate metabolic process | 5 out of 152 total entries |
| lipid metabolic process | 3 out of 298 total entries | response to chemical | 4 out of 413 total entries |
| mitochondrial translation | 3 out of 172 total entries | nucleobase-containing small molecule metabolic process | 4 out of 178 total entries |
| monocarboxylic acid metabolic process | 3 out of 140 total entries | protein folding | 4 out of 101 total entries |
| proteolysis involved in protein catabolic process | 2 out of 224 total entries | ribosomal subunit export from nucleus | 4 out of 67 total entries |
| DNA repair | 2 out of 255 total entries | mitochondrial translation | 4 out of 172 total entries |
| response to chemical | 2 out of 413 total entries | monocarboxylic acid metabolic process | 3 out of 140 total entries |
| nucleobase-containing small molecule metabolic process | 2 out of 178 total entries | transmembrane transport | 3 out of 277 total entries |

**Table S4.** Comparison between the 3D growth media and prior experimental data characterizing mucosal samples.

| Material property | 3D growth matrix | Mucosal samples | References |
| --- | --- | --- | --- |
| Stiffness (elastic modulus) | Ultra-low confinement;<br>$G' \sim 3$ Pa<br>Low confinement, $G' \sim 30$ Pa<br>High confinement, $G' \sim 350$ Pa | Soft mucus,<br>$G' \sim 10$ Pa<br>Stiff mucus,<br>$G' \sim 100$ Pa | 3D matrix<sup>6-8</sup><br>Mucosal samples<sup>9-11</sup> |
| Porosity (nano-scale) | Mesh sizes within randomly cross-linked polymer chains<br>$\sim 40$-$100$ nm | Nanometer-scale pores in human cervicovaginal mucus<br>average pore size<br>$\sim 340 \pm 70$ nm | 3D matrix<sup>6-8</sup><br>Mucosal samples<sup>12</sup> |
| Porosity (micron-scale) | Inter-particle pore spaces within the 3D matrix<br>Low confinement ($\sim 5$ microns)<br>High confinement ($\sim 2$ microns) | Pores in pig jejunum, pig ileum, and human ileum<br>Variably sized, micron-scale pores | 3D matrix<sup>6-8</sup><br>Mucosal samples<sup>13</sup> |
| Charge density | Negatively-charged polymer (Carbopol 980)<br>Highly negatively-charged polymer (ETD 2020) | Mucins carry negative charges | 3D matrix<sup>6-8</sup><br>Mucosal samples<sup>14,15</sup> |

